# Gustatory Thalamic Neurons Mediate Aversive Behaviors

**DOI:** 10.1101/2024.07.29.605702

**Authors:** Feng Cao, Sekun Park, Jordan L. Pauli, Eden Y. Seo, Richard D. Palmiter

**Affiliations:** Howard Hughes Medical Institute, University of Washington, Seattle, WA 98195, USA; Departments of Biochemistry, University of Washington, Seattle, WA 98195, USA; Department of Genome Science, University of Washington, Seattle WA 98195 USA

## Abstract

The parvicellular part of the ventral posteromedial nucleus (VPMpc) of the thalamus, also known as the gustatory thalamus, receives input from the parabrachial nucleus and relays taste sensation to the gustatory (or insular) cortex. Prior research has focussed on the role of the VPMpc in relaying taste signals. Here we provide evidence showing that VPMpc also mediates aversive behaviors. By recording calcium transients *in vivo* from single neurons in mice, we show that neurons expressing cholecystokinin and the mu-opioid receptor in the VPMpc respond to various noxious stimuli and fear memory. Chemogenetic and optogenetic activation of these neurons enhances the response to aversive stimuli, whereas silencing them attenuates aversive behaviors. The VPMpc neurons directly innervate neurons in the insular cortex and rostral lateral amygdala. This study expands the role of the VPMpc to include mediating aversive and threating signals to the insular cortex and lateral amygdala.

## INTRODUCTION

The thalamus is considered as one of the most highly interconnected brain regions, acting as a hub for relaying sensory information between different subcortical areas and the cerebral cortex^1–3^. It is composed of different thalamic nuclei and segmented into several subregions based on anatomical location and thalamic-cortical connectivity patterns^3–5^. Each subregion serves an important role, ranging from relaying sensory and motor stimulations to regulation of sleep, consciousness, and learning and memory^2–5^.

The parvicellular part of ventral posteromedial nucleus (VPMpc), also called the gustatory thalamus, is located medial to the ventral posteromedial nucleus (VPM) and contains neurons that process and transmit taste information to the insular cortex (IC)^6–8^. In the rodent gustatory sensory pathway, taste-related information from the tongue travels through cranial nerves VII, IX and X, converges in the nucleus solitarius (NTS) in the medulla, sends ascending neural projections to the parabrachial nucleus (PBN) in the dorsal pons, which projects to the VPMpc, and then to the gustatory part of the IC^6,9–13^. Neurons in VPMpc reliably encode the physiochemical identity of gustatory stimuli from different tastants delivered to the tongue and exert a strong influence on IC activity^7,14,15^. A few early studies found that VPMpc neurons also respond to tactile and thermal stimuli on the tongue of anesthetized rats^8,16^. Neurons in VPMpc participate in taste expectations; lesions of the VPMpc in rats disrupted aversive and appetitive anticipatory taste^17^, while an expected anticipatory cue improved taste processing by VPMpc neurons^18^.

Although the VPMpc is widely associated with gustatory function, little is known about its involvement in other neurological functions or innate behaviors. The VPMpc receives inputs from several different neuron clusters in PBN including *Satb2*-expressing neurons (encoding special AT-rich sequence-binding protein 2) in the lateral and waist regions of the PBN and *Calca*-expressing neurons (encoding calcitonin gene-related peptide) in the external lateral PBN^19,20^. The *Satb2* neurons have been shown to relay taste information to the VPMpc, but they project to many additional brain regions that are not known to be involved in taste processing^20,21^. The CGRP^PBN^ neurons are activated by most sensory modalities (e.g., visceral malaise, taste, temperature, pain, itch) and function as a general alarm^22–24^. Photoactivation of the CGRP^PBN^ axon terminals in VPMpc elicited freezing, fear memory and avoidance behavior, while photoinhibition of the VPMpc terminals alleviated fear response and fear memory^25^.

Among all the brain regions that receive sensory input from CGRP^PBN^ neurons, VPMpc exhibited significantly greater excitation from terminal activation than any other recorded projection sites^25^. Recent studies also found that the gustatory cortex encodes aversive taste memory and aversive states^26–28^. Given the interconnectivity of VPMpc with PBN and IC, we hypothesized that VPMpc also transmits aversive signals to the IC and mediates aversive behaviors.

Here we provide anatomical and functional evidence demonstrating that CGRP^PBN^ innervate molecularly defined neurons in VPMpc neurons that project axons to IC and rostral lateral amygdala (rLA). Using transgenic and viral strategies to deliver Cre-dependent genes to VPMpc neurons, we found that VPMpc neurons responded to multiple aversive stimuli and cues that predicted them. Together with the gain-of-function and loss-of-function studies, we demonstrate that VPMpc neurons bidirectionally contribute to behavioral responses to aversive events, providing evidence that VPMpc neurons process non-gustatory signals.

## RESULTS

### Cholecystokinin (CCK) and mu opioid receptor (OPRM1) are co-expressed in VPMpc neurons that are directly innervated by CGRP^PBN^ neurons

Prior experiments revealed extensive axonal projections to the VPMpc by CGRP^PBN^ neurons labeled with fluorescent probes and photoactivation of axon terminals in the VPMpc elicited significant biological effects^19,25^. To gain direct access to the VPMpc neurons that are targets of the CGRP^PBN^ neurons, we sought to identify useful Cre-driver lines of mice. We initially checked Cre-driver lines of mice for receptors of the neuropeptides made by CGRP^PBN^ neurons (CGRP, NTS, PACAP and TAC1) by injecting AAV carrying Cre-dependent fluorescent proteins into the VPMpc of *Calcrl^Cre^, Ntsr1^Cre^, Adcyap1r1^Cre^* or *Tacr1^Cre^* mice, respectively (Extended data Fig. 1a- d); however, none of these Cre-driver lines expressed fluorescent protein within the VPMpc (Extended data Fig. 1e-h). We then turned to Allen Mouse Brain Altas^29^ to look for candidate genes expressed in the VPMpc for which Cre-driver lines of mice existed. We focussed on two gene candidates: *Cck* and *Oprm1*. To verify their expression in VPMpc, AAV1-DIO-YFP virus was injected into VPMpc of *Cck^Cre^* or *Oprm1^Cre^* mice (Fig. 1a); YFP was expressed in the VPMpc of both lines of mice (Fig. 1b).

**Fig. 1.**
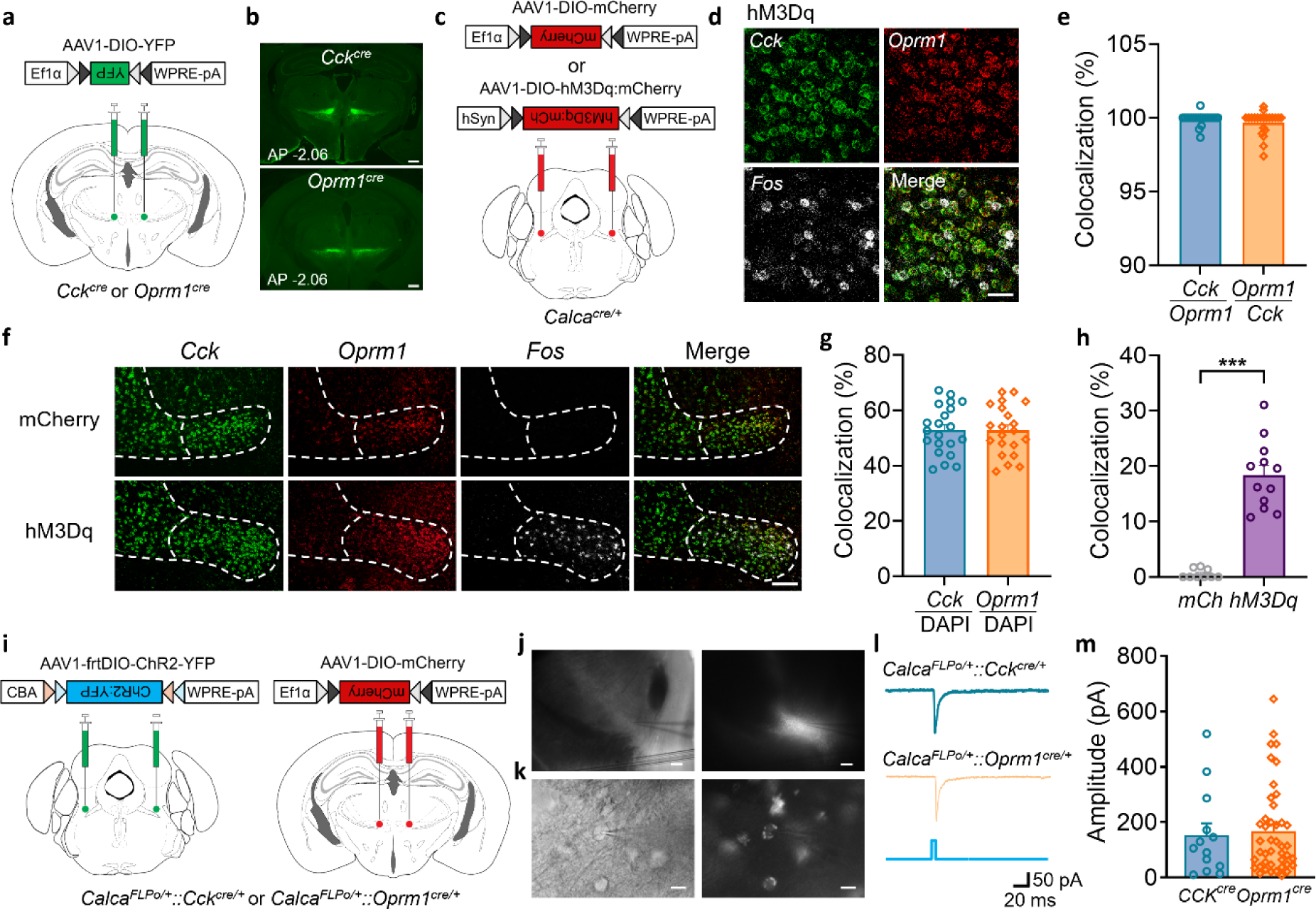
**CCK^VPMpc^ neurons and OPRM1^VPMpc^ neurons are direct innervated with CGRP^PBN^ neurons a**, Scheme showing bilateral injections of AAV1-DIO-YFP into the VPMpc of *Cck^Cre^* and *Oprm1^Cre^* mice. **b**, AAV1-DIO-YFP expression in VPMpc of *Cck^Cre^* and *Oprm1^Cre^* mice, scale bar 500 μm. **c**, Scheme showing bilateral injections of AAV1-DIO-mCherry or AAV1-DIO-hM3Dq:mCherry into the PBN of *Calca^Cre^* mice. **d**, High magnification RNAscope images showing the overlap of *Cck*, *Oprm1*, *Fos* in VPMpc of mice injected AAV1-DIO-hM3Dq:mCherry into PBN, scale bar 40 μm. **e**, Summary data of the *Cck* and *Oprm1* colocalization. **f**, Low magnification images showing RNAscope of *Cck*, *Oprm1,* and *Fos* in VPMpc of mice injected AAV1-DIO-mCherry and AAV1-DIO- hM3Dq:mCherry into PBN, scale bar 150 μm. **g**, Summary data showing the percentage of DAPI- positive cells that express *Oprm1* or *Cck*. **h**, Summary data of *Fos* and *Cck* colocalization in both mCherry and hM3Dq groups. **i**, Scheme showing bilateral injections of AAV1-frtDIO-ChR2:YFP or AAV1-DIO-mCherry into the PBN and VPMpc of *Calca^FLPo^::Cck^Cre^* and *Calca^FLPo^::Oprm1^Cre^*mice. **j**, VPMpc under 4× differential interference contrast (DIC) objective (left) and fluorescence object (right) images with recording electrode on top of it, scale bar 150 μm. **k**, Image of VPMpc neurons under 40× DIC objective (left) and fluorescence object (right) images with recording electrode attach on the recording neuron, scale bar 10 um. **l**, Sample traces and **m**, Summary figure of 455-nm blue-light pulse of light-evoked EPSCs in the VPMpc neurons of *Calca^FLPo^::Cck^Cre^*and *Calca^FLPo^::Oprm1^Cre^* mice. Circled individual data points represent *Cck* or *Cck^Cre^* mice, diamonds represent *Oprm1* or *Oprm1^Cre^* mice in the summarized bar graphs. Data are represented as mean ± SEM. ***P < 0.001.

To determine whether CCK^VPMpc^ or OPRM1^VPMpc^ neurons are downstream of CGRP^PBN^ neurons, we injected AAV carrying Cre-dependent hM3Dq:mCherry, an excitatory DREADD (Designer Receptors Activated Only by Designer Drugs)^30,31^ or just mCherry as control in the PBN of *Calca^Cre^* mice (Fig. 1c). After allowing 6 weeks for viral expression, hM3Dq was activated by injection of its ligand, clozapine-N-oxide (CNO, 1 mg/kg, i.p.) and the brains were prepared 45 min later for *in situ* hybridization using RNAscope probes for *Cck, Oprm1* and *Fos*. *Cck* mRNA was colocalized with *Oprm1* mRNA in nearly all VPMpc neurons (Fig. 1d-f), which were about 50% of all DAPI-positive cells in VPMpc (Fig. 1h). After CNO administration, *Fos* was expressed in >18% of either the *Cck-* or *Oprm1*-expressing neurons in mice with hM3Dq:mCherry virus and given CNO, but in <0.6% of cells when CNO was injected in mice with the control mCherry virus (Fig. 1f, h). The percentage of *Fos* mRNA colocalized with either *Cck* or *Oprm1* mRNA in VPMpc was >98%. We conclude that *Cck* and *Oprm1* are co-expressed in the same neurons, and they express *Fos* mRNA after activation of CGRP^PBN^ neurons with hM3Dq and CNO.

We used electrophysiology and optogenetic methods to determine whether CCK^VPMpc^ and OPRM1^VPMpc^ neurons are directly innervated by the ChR2-expressing presynaptic terminals from CGRP^PBN^ neurons. To specifically target the CCK^VPMpc^ and OPRM1^VPMpc^ neurons, we generated mice expressing *Calca^FLPo^ ::Cck^Cre^* or *Calca^FLPo^:: Oprm1^Cre^* and then injected AAV carrying FLPo-dependent ChR2-YFP into PBN and Cre-dependent mCherry into VPMpc (Fig. 1i). After allowing 6 weeks for viral expression, we prepared brain slices and recorded excitatory post-synaptic currents (EPSCs) in response to photoactivation in CCK^VPMpc^ or OPRM1^VPMpc^ neurons (i.e., mCherry-positive soma with nearby YFP fibers from Calca neurons, Fig. 1j, k).

Blue-light, optical stimulation of CGRP^PBN^ terminals in the VPMpc evoked EPSCs in both CCK^VPMpc^ (13/13) and OPRM1^VPMpc^ (41/43) neurons of *Calca^FLPo^::Cck^Cre^*and *Calca^FLPo^::Oprm1^Cre^* mice, with a <3-ms onset latency (Fig. 1l). There was no significant difference between the evoked EPSC amplitude in CCK^VPMpc^ and OPRM1^VPMpc^ neurons (Fig. 1m). Bath application of α-amino-3- hydroxy-5-methyl-4-isoxazolepropionic (AMPA) receptor antagonist NBQX (10 μM) and N-methyl-D-aspartate (NMDA) receptor antagonist (2R)-amino-5-phosphonovaleric acid (APV, 50 μM) to the brain slices completely blocked the EPSCs (Extended Data Fig. 2a, b). We also investigated blue-light induced synaptic release from isolated ChR2-transduced PBN axons terminals in the VPMpc. Bath application of the Na^+^ channel blocker, tetrodotoxin (TTX, 1 μM), abolished light-evoked EPSCs which could be reinstated by the subsequent addition of 1 mM K^+^ channel blocker 4-aminopyridine (4AP) (Extended Data Fig. 2c-e), although the onset latency was typically delayed^32,33^. These results indicate that the light-evoked EPSCs in CCK^VPMpc^ or OPRM1^VPMpc^ are mono-synaptic, glutamatergic and action-potential dependent. Thus, we conclude that *Cck* and *Oprm1* genes are co-expressed in VPMpc neurons, and these neurons receive direct, excitatory input from CGRP^PBN^ neurons.

### VPMpc neurons respond to aversive sensory modalities

CGRP^PBN^ neurons serve as a general alarm and are activated by many aversive events and cues that predict them^22,24^. Because CGRP^PBN^ neurons innervate CCK and OPRM1 neurons in VPMpc, we hypothesized that they would also respond to aversive stimuli. We used *Cck^Cre^* mice and 1- photon calcium imaging to evaluate whether VPMpc neurons respond to aversive stimuli. *Cck^Cre^* mice were injected with AAV carrying a Cre-dependent calcium indicator, GCaMP6m; then a gradient refractive index (GRIN) lens was implanted over the VPMpc to monitor the CCK^VPMpc^ neuron activity in freely moving mice (Fig. 2a). We examined the responses to a 90-dB hand clapping sound (Fig. 2b), a 1-s air puff onto the head (Fig. 2e), a 1-s tail pinch (Fig. 2h), or 10-s of lifting the mice out of the cage by their tail (Fig 2k). We applied these different sensory modalities to mice individually and recorded the calcium signal before and after each stimulus (78 neurons from 4 mice). The 90-dB hand clapping sound generated a smaller increase in calcium activity (Fig. 2b-d) than either the air puff (Fig. 2 e-g) or tail pinch (Fig. 2h-j). Lifting the mouse by the tail also elevated the calcium activity with a gradual increase during the 10-s lifting period, which remained elevated for 10 s after the mice were returned to the cage (Fig. 2k-m).

**Fig. 2.**
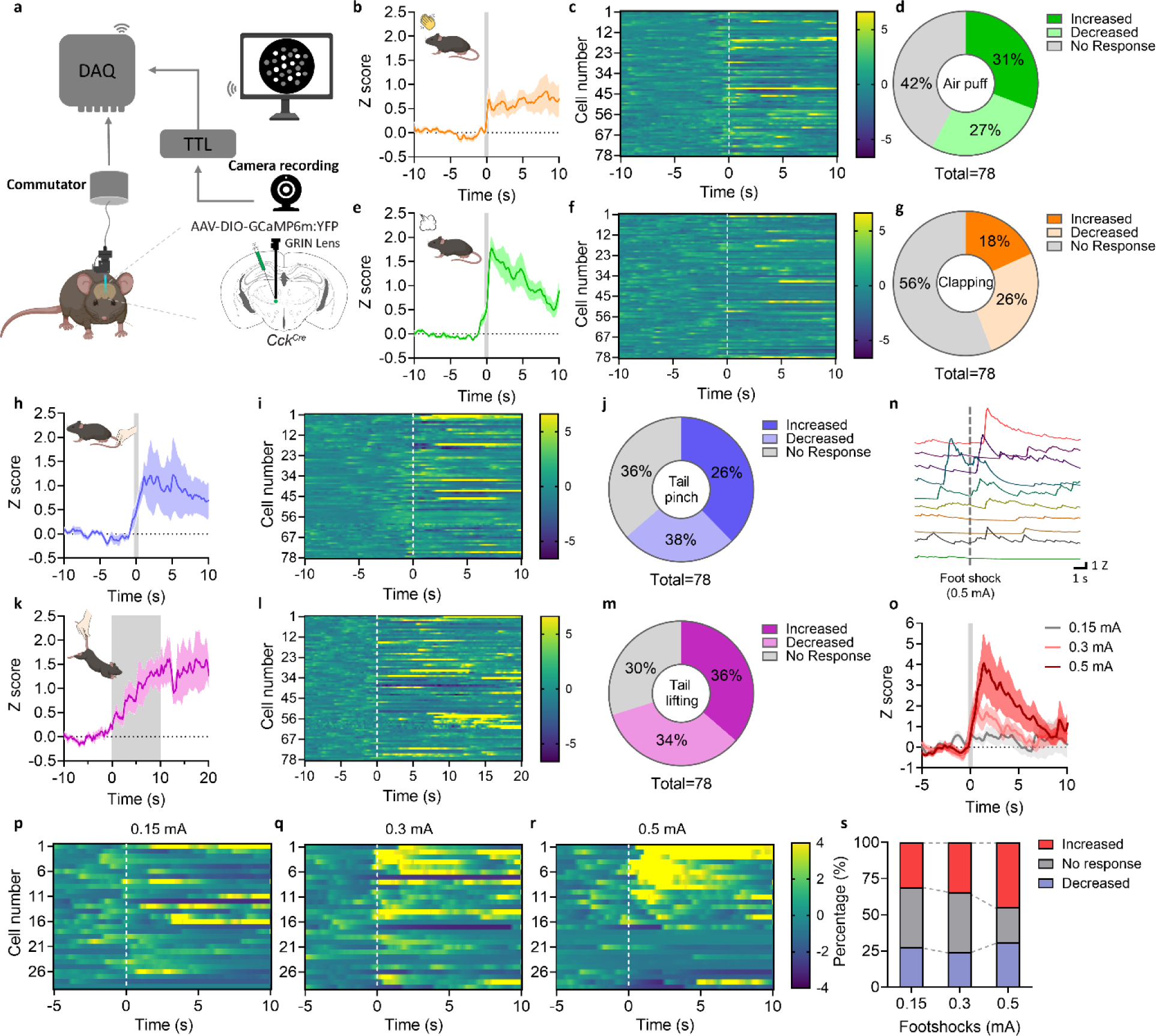
VPMpc neurons respond to aversive sensory stimuli **a**, Scheme of 1-photon single-cell calcium imaging in freely moving mice. **b-m**, Average traces with schematic diagram (**b, e, h, k**), heat maps of individual neuronal responses (**c, f, i, l**), percentage of increased, decreased, and no response CCK^VPMpc^ neurons (**d, g, j, m**) in response to hand clapping sound, air puff, tail pinch, and tail lifting. Vertical dash lines: onset of each sensory stimulus. **n**, Sample calcium fluorescence activity trace of 10 CCK^VPMpc^ neurons when mice underwent 0.5-mA foot shock. **o**, Average traces, **p-r**, heat maps of CCK^VPMpc^ neuron calcium fluorescence activity in response to 0.15, 0.3 and 0.5 mA foot shocks. Vertical dash lines: onset of the foot shocks **s**, Percentage of increased, decreased, and no response CCK^VPMpc^ neurons in response to 0.15, 0.3, and 0.5 mA foot shocks.

We applied three different shock intensities (0.15, 0.3 and 0.5 mA) to mice placed on a shock grid and recorded the calcium activity generated by each intensity. All shock intensities generated increases in calcium activity that were positively correlated with shock intensities (Fig. 2n-s). The 0.5-mA shock activated the largest number of neurons (Fig. 2s), and most of them were also activated by the 0.3-mA shock (Fig. 2q, r). We conclude that VPMpc neurons respond to many aversive modalities.

### Fear memory activates VPMpc neurons

We analyzed CCK^VPMpc^ neuronal activity during auditory fear conditioning, an associative learning paradigm in which a tone, a conditioned stimulus (CS), was paired with a foot shock, an unconditioned stimulus (US). We used the same mice with GCaMP6m expression and GRIN lens implantation as described above and measured the calcium dynamics of CCK^VPMpc^ neurons while performing a three-day fear conditioning protocol (Fig. 3a-c). On Day 1, only a few CCK^VPMpc^ neurons were activated while mice were exposed to 10 trials of a 10-s tone (Fig. 3d, g, Extended Data Fig. 3). On Day 2, with 10 trials of the CS-US association, many CCK^VPMpc^ neurons were activated by the foot shock (Fig. 3e, h, Extended Data Fig. 4), which is consistent with the calcium activity change under foot shock in previous experiment (Fig. 2o). On Day 3, the mice were exposed to the tone alone in a novel context; many CCK^VPMpc^ neurons responded during the 10-s tone, and the activity persisted beyond the 10 s of recording following the tone (Fig. 3f, I, Extended Data Fig. 5); the response was much greater than the calcium activity on Day 1.

**Fig. 3.**
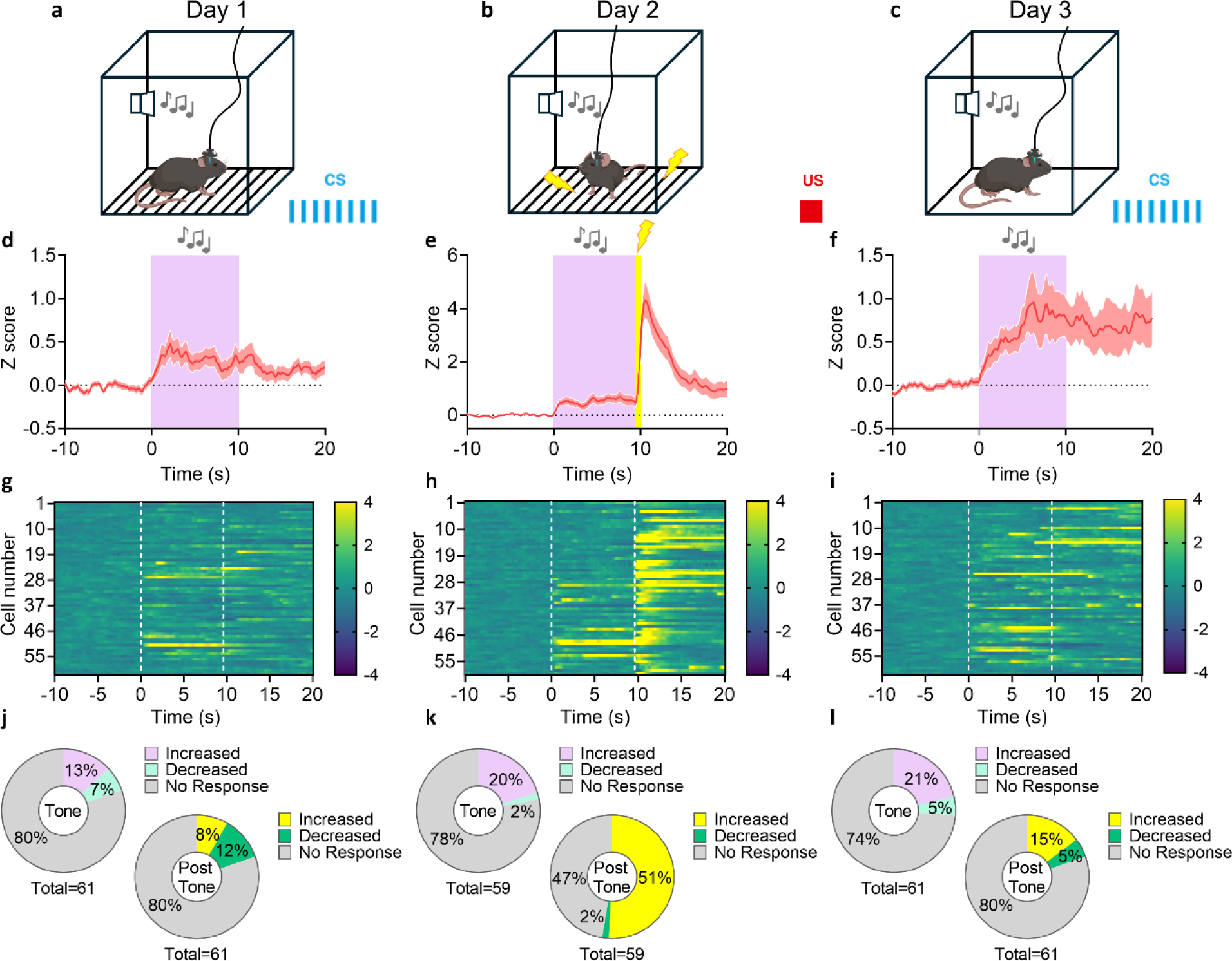
VPMpc neurons contribute to fear-learning memory recall **a-c**, Schematic diagram of calcium imaging in 3-day fear conditioning paradigm. **d-i**, Average traces (**d, e, f**) and heat maps of individual neuronal responses (**g, h, i**) of CCK^VPMpc^ neuron calcium fluorescence activity 10 s before tone, 10 s during tone, 10 s after tone on Day 1, Day 2 and Day 3. Tone starts at 0 s in all traces, shock starts at 9.5 s on Day 2. Vertical dash lines: onset of each tone and foot shock. **j-l**, Percentage of increased, decreased and no response CCK^VPMpc^ neurons during 10-s tone (left) and 10-s post-tone (right) on Day 1 (**j**), Day 2 (**k**) and Day 3 (**l**).

Compared to the activity change during Day 1 (Fig. 3j), the proportion cells responding on both Day 2 (Fig. 3k) and Day 3 (Fig. 3l) was significantly increased. These calcium-imaging studies reveal that CCK^VPMpc^ neurons become activated by the CS only after conditioning and reflect the recall of the fear memory, like that observed in CGRP^PBN^ neurons^22^.

### Inactivation of CCK/OPRM1^VPMpc^ neurons attenuates the response to aversive stimuli

To determine whether CCK/OPRM1^VPMpc^ neurons are necessary for the aversive behavioral responses, we silenced them with tetanus toxin (TetTox), which cleaves synaptic vesicle protein synaptobrevin, thereby preventing neurotransmitter release and signaling to post-synaptic neurons^34,35^. *Oprm1^Cre^* or *Cck^Cre^* mice were bilaterally injected in the VPMpc with AAV carrying Cre-dependent tetanus toxin (AAV_DJ_-EF1a-DIO-GFP:TetTox) or YFP (AAV1-DIO-YFP) as control (Fig. 4a). We first examined tactile sensitivity to a set of von Frey filaments with different stiffness. As shown in Fig. 4b, mice injected with TetTox virus manifested a significant increase in the paw-withdraw threshold response measured by von Frey filaments, indicating that silencing them decreases tactile sensitivity. We also observed that mice with TetTox had fewer nocifensive behaviors and diminished escape attempts during exposure to noxious heat compared to control mice with only YFP (Fig. 4c, d).

**Fig. 4.**
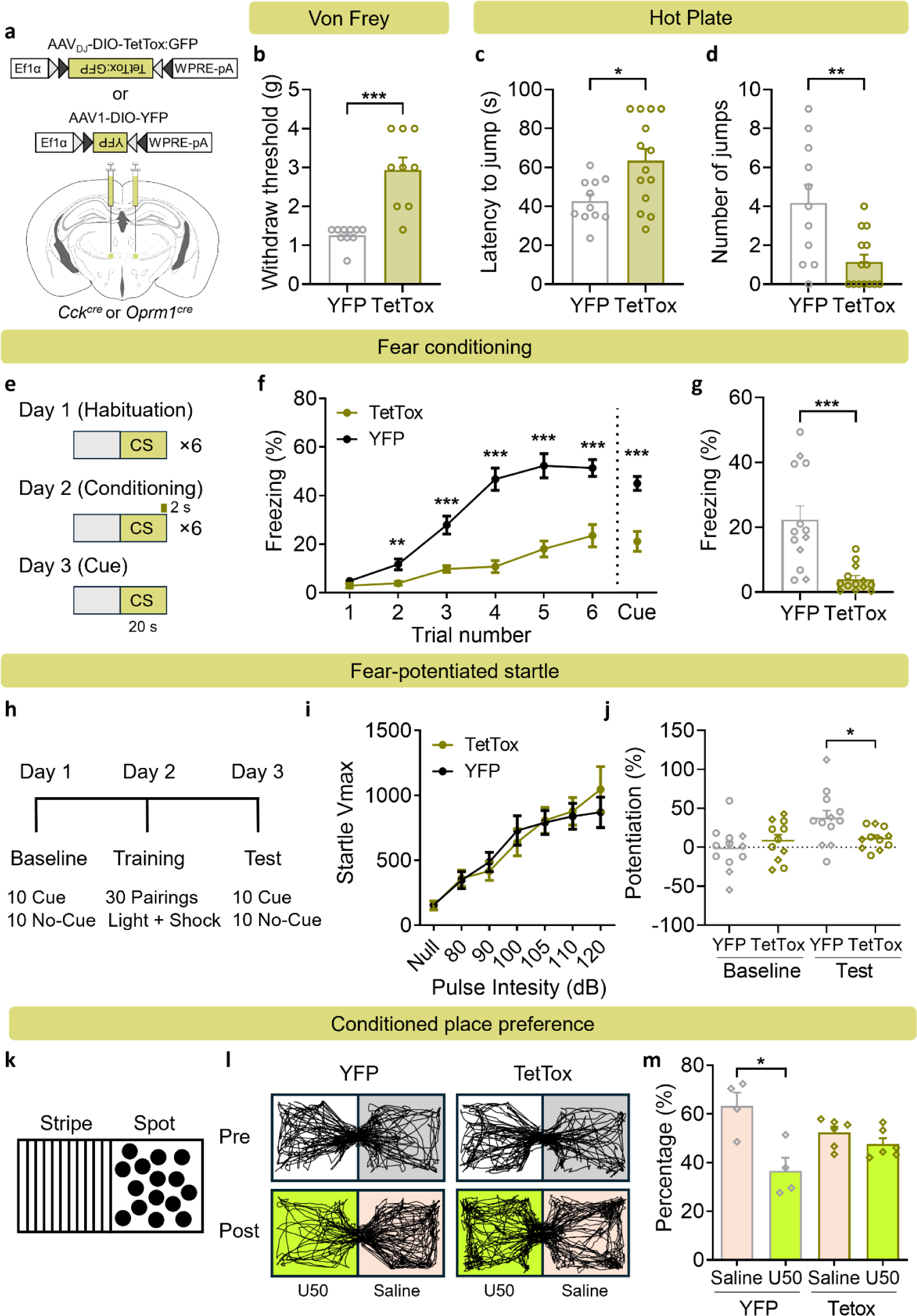
Inactivation of CCK/OPRM1^VPMpc^ neurons attenuates aversive responses **a**, Scheme showing bilateral injections of AAV_DJ_-DIO-TetTox-GFP and AAV1-DIO-GFP into the VPMpc of *Cck^Cre^* and *Oprm1^Cre^* mice. **b**, Von Frey test showing increased withdraw threshold in TetTox injected mice. **c, d**, Hot plate test showing increased latency to first jump (**c**) and decreased number of jumps (**d**) in TetTox-expressing mice. **e**, Experimental paradigm for 3-day cue-dependent fear conditioning test. **f,** Summary data showing decreased freezing responses of TetTox-expressing mice during training and cue test 24 h after conditioning (n = 13 per group, two-way repeated measures ANOVA, F (1, 24) = 71.60, P < 0.0001). **g**, Summary data showing decreased freezing response of TetTox-expressing mice in context test 24 h after conditioning. **h**, Experimental paradigm for 3-day fear-potentiated startle test. **i**, Startle test showing no difference between startle responses to multiple pulse intensities in YFP and TetTox-expressing mice (YFP: n = 12, TetTox: n = 14; two-way repeated measures ANOVA, F (1, 24) = 0.0122, P = 0.9128). **j**, Fear-potentiated startle test showing higher conditioned startle response after light cue training in YFP mice but not in TetTox mice. **k**, Scheme showing conditioned place preference chamber with stripes on one side and spots on the other. **l**, Representative mobility traces showing no preference before U50-paired conditioning (Day 1) in YFP and TetTox mice, higher preference in Saline chamber after U50 paired conditioning (day 4) in YFP but not TetTox mice. **m**, Summary data showing aversion to U50 in YFP mice but not TetTox mice after U50-paired conditioning. Circled individual data points represent *Cck^Cre^* mice, diamonds represent *Oprm1^Cre^*mice in the summarized bar graphs. Data are represented as mean ± SEM. *P < 0.05, **P < 0.01, ***P < 0.001.

We also tested whether inhibiting these neurons affected fear memory formation in the 3-day fear conditioning paradigm described above (Fig. 4e). Mice injected with the control virus had an increased freezing response to the shock during the training trials, whereas the response of the TetTox-expressing mice was much weaker (Fig. 4f); the response to the tone alone in a novel box the next day was also weaker (Fig. 4f). The control (YFP) mice had a strong freezing response to the original box when tested the next day, but the TetTox- expressing mice did not (Fig. 4g). We also examined the response of the TetTox mice in a 3-day fear-potentiated startle test, in which a light predicted a foot shock on Day 2 and we measured the startle response to the light on Day 3 (Fig. 4h). The startle responses to a range of pulse intensities were similar between the two groups of mice before training; however, after training, the light (CS cue) enhanced the startle response in the YFP (control) group but not in the TetTox group (Fig. 4j). These results confirm that the TetTox- expressing mice have difficulty learning or remembering the CS-US association.

To ascertain the affective state generated by silencing CCK/OPRM1^VPMpc^ neurons, we used a conditioned place aversion (CPA) test with two chambers that had either stripes or spots on the wall (Fig. 4k). On the pre-test day, both YFP and TetTox-expressing mice exhibited equal preference towards the two chambers (Fig. 4l, m). After pairing one chamber with U50488 (U50, 5 mg/kg), a kappa opioid receptor agonist that is aversive^36,37^, mice injected with YFP demonstrated an aversion to U50, whereas mice injected with TetTox did not (Fig. 4l, m). Taken together, these results reveal that inactivation of CCK/OPRM1^VPMpc^ neurons interfered with the normal responses to aversive stimuli.

### Activation of CCK/OPRM1^VPMpc^ neurons enhances the response to aversive stimuli

We next examined whether activation of CCK/OPRM1^VPMpc^ neurons could alter the response to aversive stimuli and aversive memory. We bilaterally injected AAV carrying Cre-dependent hM3Dq:mCherry (or mCherry as control) into VPMpc of *Oprm1^Cre^*or *Cck^Cre^* mice (Fig. 5a). The effect of hM3Dq activation by CNO was verified in brain slices using cell-attached voltage clamp to measure spontaneous cell firing before and after CNO application. Bath application of CNO increased spontaneous cell firing in neurons with hM3Dq:mCherry expression, but not neurons with only mCherry expression (Fig. 5b). We examined whether activation of CCK/OPRM1^VPMpc^ neurons changed tactile sensitivity to von Frey filaments of different stiffness. The hM3Dq- expressing mice treated with CNO had a significant decrease in the paw-withdraw threshold (Fig. 5c), indicating that activating these neurons induced allodynia. We also found that mice with hM3Dq:mCherry had more nocifensive responses and increased escape attempts during exposure to noxious heat compared with mice with mCherry (Fig. 5d-f). These results indicate that chemogenetic activation of CCK/OPRM1^VPMpc^ neurons enhances the nocifensive response to noxious stimuli.

**Fig. 5.**
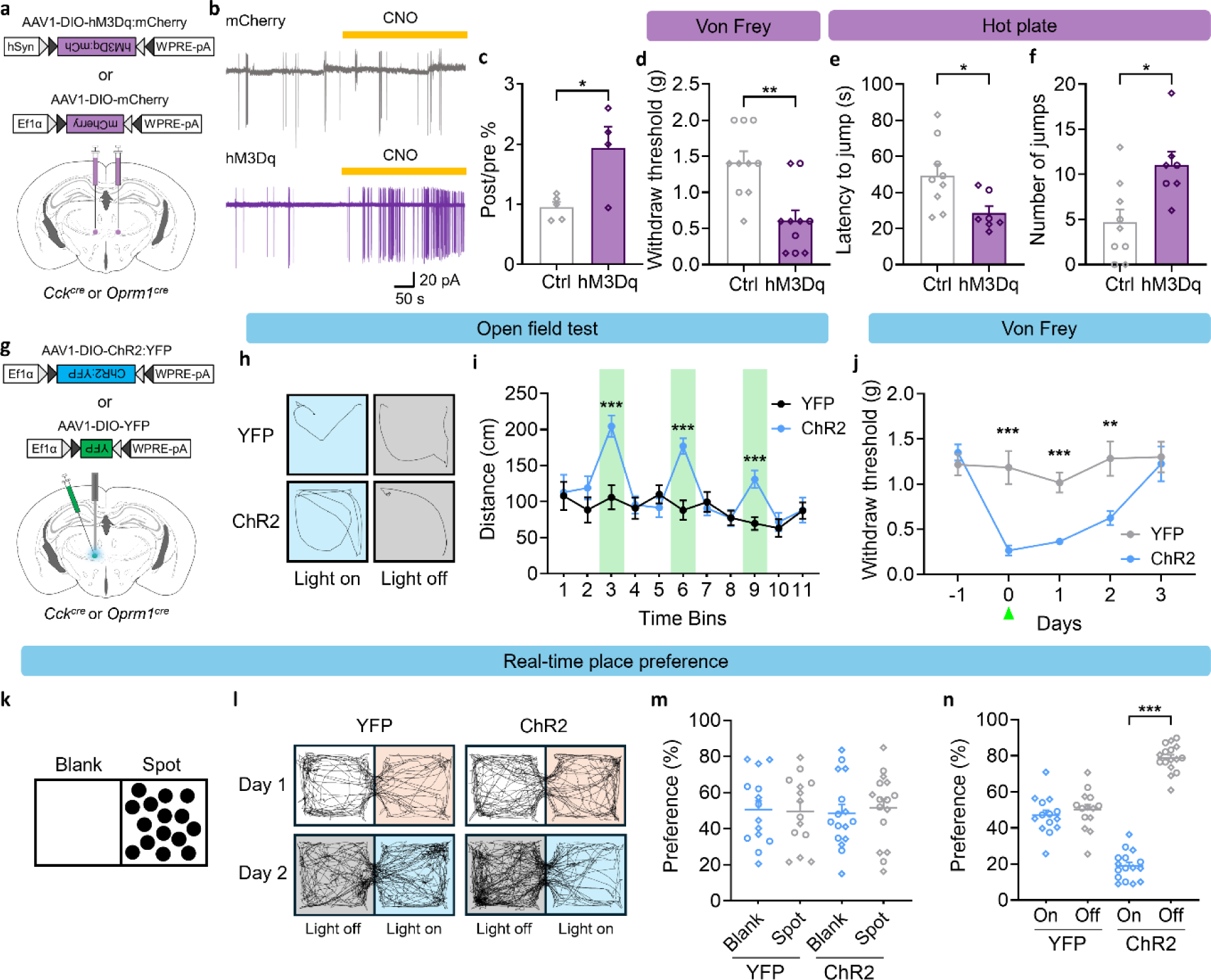
Activation of CCK/OPRM1^VPMpc^ neurons enhances aversive responses **a**, Scheme showing bilateral injections of AAV1-DIO-hM3Dq:mCherry and AAV1-DIO-mCherry into the VPMpc of *Cck^Cre^* and *Oprm1^Cre^* mice. **b**, Sample traces and **c**, summary figure showing CNO-induced spontaneous cell firing in hM3Dq but not mCherry-injected mice. **d**, Von Frey test showing decreased paw withdraw threshold in hM3Dq-injected mice. **e, f**, Hot plate test showing decreased latency to jump (**e**) and increased number of jumps (**f**) in hM3Dq-injected mice. **g**, Scheme showing unilateral injections of AAV1-DIO-ChR2:YFP and AAV1-DIO-YFP into the VPMpc of *Cck^Cre^* and *Oprm1^Cre^* mice. h, Representative mobility traces of mice with YFP and ChR2 during light on (bin 3) and light off (bin 4). **i**, Open field test showing increased distance traveled in photo-stimulated (bin 3, 6, 9) ChR2 mice (n = 10 per group, two-way repeated measures ANOVA, F (1, 18) = 11.45, P = 0.0033). **j**, Von Frey test showing continuous days of decreased paw-withdraw threshold after 5 min of photo-stimulation (10 ms, 10 Hz, 2 mW) with ChR2-expressing mice (YFP: n = 6, TetTox: n = 8; two-way repeated measures ANOVA, F (1, 12) = 13.5, P = 0.0032). **k**, Scheme of RTPP paradigm. **l**, Representative mobility traces showing no side preference in both YFP and ChR2 mice without photostimulation on Day 1, higher preference in light-off chamber in ChR2-expressing mice but not in YFP mice on Day 2. **m**, YFP mice and ChR2 mice had similar side preference without optical stimulation. **n**, Photo activation of ChR2- injected mice led to avoidance of light-paired side. Circles represent *Cck^Cre^* mice, diamonds represent *Oprm1^Cre^*mice in the summarized bar graphs. Data are represented as mean ± SEM. *P < 0.05, **P < 0.01, ***P < 0.001.

To further examine the effect of activating CCK/OPRM1^VPMpc^ neurons, we unilaterally injected AAV carrying Cre-dependent channelrhodopsin, ChR2:YFP (or YFP as control) into VPMpc of *Cck^cre^* and *Oprm1^Cre^* mice and fiber-optic cannulae were placed over the VPMpc (Fig. 5g). We placed both groups of mice in an open-field chamber and recorded their mobility throughout eleven 30-s time bins with 473-nm blue light stimulation (10 Hz, 10 ms, 2 mW) during time bins 3, 6 and 9 (Fig. 5h). Mice expressing ChR2 exhibited higher travel distance in each light-stimulated bin, whereas control mice had the same travel distance across all bins (Fig. 5h, i). We also examined the tactile sensitivity in both control and ChR2 groups after they received 5-min of photo-stimulation (10 Hz, 10 ms, 2 mW). Photoactivating ChR2-expressing CCK/OPRM1^VPMpc^ neurons had the same allodynia effect as activating hM3Dq-expressing VPMpc neurons with CNO (Fig. 5d, j). Interestingly, the allodynia caused by photo-stimulation lasted 2 days (Fig. 5j), indicating that brief activation of CCK/OPRM1^VPMpc^ neurons can trigger prolonged tactile sensitivity.

To determine the affective state when activating CCK/OPRM1^VPMpc^ neurons, we used a real-time place preference (RTPP) test to ask whether mice prefer or avoid optical stimulation.

A two-chambered box with one side blank and the other decorated with spots was used for these experiments (Fig. 5k). The mice could freely explore the two chambers for 10 min on Day 1. Both groups of mice demonstrated similar preferences towards the two chambers without optical stimulation on Day 1 (Fig. 5l, m). We then paired optical stimulation (10 Hz, 10 ms, 2 mW) in only one of the two chambers. Mice with only YFP expression had no preference for either side, whereas mice with ChR2 stimulation robustly avoided the side with optical stimulation (Fig. 5l, m), indicating that photoactivation of CCK/OPRM1^VPMpc^ neurons is aversive. We chose these relatively mild stimulation conditions because more robust stimulation (30 Hz, 10 ms, 10 mW) resulted in rotation behavior and body distortion (Supplementary Videos 1 & 2). Taken together, these results demonstrate that both chemogenetic and optogenetic activation of CCK/OPRM1^VPMpc^ neurons enhances the responses to aversive stimuli.

### Axonal projections of CCK^VPMpc^ neurons

VPMpc neurons have long been known to transmit taste signals to the forebrain across different species in mammals^6,8,38,39^. The projection from the VPMpc to the IC is a major component of the gustatory pathway from the taste buds in the tongue to the gustatory cortex^2,7,39^. To identify the axonal projections from VPMpc, we injected AAV expressing Cre-dependent fluorescent proteins (AAV-DIO-YFP and AAV-DIO-synaptophysin:mCherry) into the VPMpc of *Cck^cre^*mice. A small volume of virus (15 nl) was injected and only those mice with expression restricted to the VPMpc were used to examine axonal projections (Fig. 6a-c). To obtain the whole brain projection pattern, we cut the entire brain into 35-μm sections and imaged every third coronal brain section. The imaged sections were stitched together and registered to view the expression of YFP and mCherry throughout the brain (Supplementary Video 3). Other than the strong projection to IC, we also observed a weaker projection to the lateral amygdala (LA) (Fig. 6d). After quantifying the signal intensity throughout the IC, we observed expression of YFP and mCherry at three collected sections (Bregma +0.75 to -0.4 mm), which covers the medial and posterior of IC (Extended Data Fig. 6 a-c). We also observed two separate bands in different cortical layers; one of them was in layer 1 and the other larger band was in layers 3-4 (Extended Data Fig. 6 b, c). The VPMpc projection to the LA (Bregma -1.3 to - 1.8 mm) was restricted to the rostral portion of the LA (Extended data Fig. 7a-c).

**Fig. 6.**
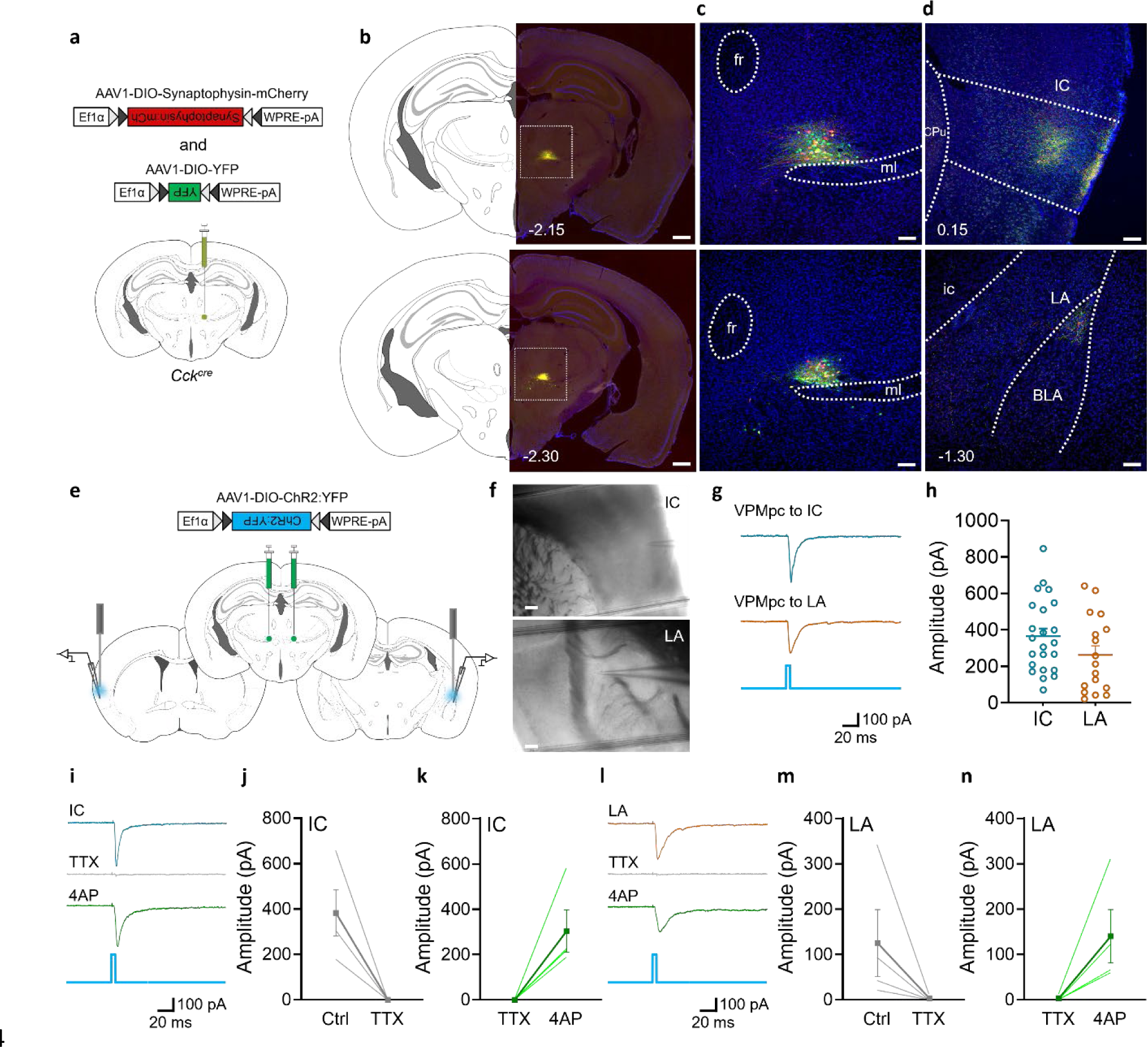
CCKVPMpc neurons send direct input to IC and LA **a**, Scheme showing unilateral injection of AAV1-DIO-mYFP and AAV1-DIO- Synaptophysin:mCherry into the VPMpc of *Cck^Cre^* mice. **b**, YFP and Synaptophysin:mCherry expression in VPMpc of *Cck^Cre^* mice, scale bar 500 μm. **c**, Sample images showing higher magnification of YFP and Synaptophysin:mCherry expression in VPMpc of *Cck^Cre^* mice, scale bar 100 μm. **d**, Sample images showing VPMpc axon terminals in IC and LA, scale bar 100 μm. See Extended Data Fig. 6 and 7 for higher magnification of IC and rostral LA. **e**, Scheme showing bilateral injections of AAV1-DIO-ChR2:YFP into the VPMpc, light-evoked electrophysiological recording in IC and LA in *Cck^Cre^* mice.**f**, Recording electrode on top of IC and LA, scale bar 150 μm. **g**, Sample traces and **h**, a summary figure of 455-nm blue-light pulse of light-evoked EPSCs in the IC and LA neurons of *Cck^Cre^* mice. **i-n**, Sample traces (**i, l**), summary figures of TTX blocked EPSCs (**j, m**) and 4AP restated EPSCs (**k, n**) in IC and LA neurons of *Cck^Cre^* mice. Data are represented as mean ± SEM.

We used electrophysiology and optogenetic methods to determine whether VPMpc neurons make functional connections with IC and rLA. We injected AAV carrying Cre-dependent ChR2 (AAV1-Ef1a-DIO-ChR2:YFP) into the VPMpc of *Cck^cre^* mice (Fig. 6e). After 6 weeks for viral expression, we prepared brain slices containing IC and rLA and recorded the activity of neurons adjacent to YFP-labeled axon terminals in these two brain regions (Fig. 6e, f). Light-evoked EPSCs were captured in both IC (23/23) and rostral LA (17/19) neurons (Fig. 6g). Although the amplitude of EPSCs in rLA were slightly smaller than those in the IC, there was no significant difference between them (Fig. 6g, h). Bath application of TTX (1 μM) blocked the EPSCs in both brain regions, which were restored by adding 4AP (1 mM) (Fig. 6i-n), suggesting both IC and LA receive direct monosynaptic inputs from CCK^VPMpc^ neurons. Thus, we conclude that CCK^VPMpc^ neurons send direct efferent projections to IC and rLA.

## DISCUSSION

We demonstrated that CCK^VPMpc^ neurons and OPRM1^VPMpc^ neurons are innervated by CGRP^PBN^ neurons. Electrophysiological recordings provided functional evidence that the input from CGRP^PBN^ neurons to CCK^VPMpc^ or OPRM1^VPMpc^ neurons is mono-synaptic and glutamatergic.

Calcium imaging in freely moving mice revealed that *Cck* neurons in the VPMpc respond to different aversive stimuli including hand clap, air puff, tail pinch, tail lifting and foot shock. They also responded to a tone (conditional stimulus) after it was associated with a foot shock.

Inactivation of CCK^VPMpc^ or OPRM1^VPMpc^ neurons produced deficits in their responses to aversive stimuli and activation of these neurons enhanced their responses to aversive stimuli. We also revealed direct axonal projections from VPMpc neurons to neurons in the IC and rLA.

Quantitative *in situ* hybridization data revealed that *Cck* and *Oprm1* are co-expressed 98% of the neurons and they represent ∼50% of all DAPI-positive cells in VPMpc. Recent validated isotropic fractionator demonstrated the ratio of neurons to glia is close to 1:1^40,41^; therefore, the other 50% are likely to be glial cells. Thus, the *Cck/Oprm1* neurons represent most, if not all, of the VPMpc neurons; single-cell RNA sequencing may reveal cell-type heterogeneity of VPMpc neurons. Because of the co-expression, we used *Cck^Cre^*and/or *Oprm1^Cre^* mice in these studies.

Electrophysiological recording from either CCK^VPMpc^ or OPRM1^VPMpc^ neurons revealed that almost all of them responded to poststimulation of ChR2-expressing axon terminals from the CGRP^PBN^ neurons with a similar mean amplitude of ∼150 pA. In retrospect, it is not surprising that CCK/OPRM1^VPMpc^ neurons respond to aversive stimuli because of the strong input from CGRP^PBN^ neurons that are known to respond to most aversive stimuli. The glutamate signaling from the CGRP^PBN^ neurons is likely to be the most relevant transmitter, because the receptors for some of neuropeptides (CGRP, NTS, PACAP and Substance P) made by CGRP^PBN^ neurons were not detected in the VPMpc. In contrast, neuropeptide signaling from CGRP^PBN^ neurons to the central nucleus of the amygdala is critical for fear learning and memory^42^.

Inactivating the CCK^VPMpc^ neurons greatly attenuated the freezing response to foot shocks both during fear learning, and the memory of the context or cue 24 hours later. These results resemble that achieved by inactivating CGRP^PBN^ neurons^23,25^. Optogenetic activation of *Calca* neurons can substitute for a foot shock in this fear-conditioning paradigm^23,25^ and the photoactivation of *Calca* neuron axonal projections in the VPMpc was able to substitute for foot shock^25^. However, directly activating the CCK^VPMpc^ neurons was unable to serve as an unconditioned stimulus in the fear conditioning paradigm (F.C., unpublished). The central tegmental fiber track from the PBN to the forebrain passes below the VPMpc^20,25^; thus, a likely explanation for this discrepancy is that photoactivation of CGRP^PBN^ neuron terminals in the VPMpc also activated fibers of passage. This result emphasizes the importance of being able to directly activate molecularly defined neurons in a brain region of interest.

Optogenetic activation of CCK^VPMpc^ neurons resulted in elevated tactile sensitivity of the hind paws with similar kinetics to that observed when activating CGRP^PBN^ neurons. Five minutes of stimulation of either the *Calca* neurons in the PBN or *Cck* neurons in the VPMpc resulted in allodynia lasting 3 days. Silencing the activity of VPMpc neurons with TetTox resulted in a sustained analgesia in the von Frey, tactile-sensitivity test. These results suggest that VPMpc neurons are part of the neural circuitry downstream of the PBN that can promote nociplastic pain – defined as pain without nerve damage^43^.

Norgren and Leonard reported in 1971 that an area in the pons now known as the PBN includes taste-responsive neurons^44^ and they reported 2 years later that the PBN neurons relay taste signals to the gustatory thalamus, i.e., the VPMpc^13^. These results have been extensively confirmed in subsequent years^9–12^. The taste-responsive neurons in the PBN were primarily located in the waist and adjacent areas with some of them interspersed within the superior cerebellar peduncle fiber tract^12,45–47^. Neurons expressing the transcription factor *Satb2* reside in this region of the PBN, they respond to all tastants, and project to the VPMpc^21,48^. *Satb2*- expressing neurons in the PBN do not overlap with *Calca* neurons^20,48^. CGRP^PBN^ neurons are activated by bitter tastes^49^ and inactivating them attenuates responses to aversive (bitter and sour) tastes^48^. Our observation that most of the neurons in the VPMpc that express *Cck* and *Oprm1* are innervated by CGRP^PBN^ neurons suggests that the synaptic inputs from *Satb2* and *Calca* neurons likely overlap in many VPMpc neurons. Indeed, we have shown using electrophysiology techniques that most of the same VPMpc neurons respond to photoactivation of axon fibers from either *Satb2* or *Calca* neurons in the PBN (to be published elsewhere). Deciphering how taste signaling by the VPMpc and the more general aversive signaling described here are distinguished in the VPMpc and downstream nuclei remains an intriguing puzzle.

VPMpc neurons have been shown respond to gustatory signals in both anesthetized and alert animals^6,8,16,18,50^. In an anesthetized rat study, VPMpc neurons also responded to thermal and tactile stimulation; applying distilled water at 0 and 37 °C into the open mouth and pinching tongue or palate by either a glass rod or non-serrated forceps could elicit VPMpc responses in anesthetized rats^8,16^. In our study, we revealed VPMpc neurons respond to tactile stimulation of the paws and exposure to a hot plate. More experiments are needed to elucidate whether the same VPMpc neurons respond to taste, thermal and tactile stimulation. We also observed behavioral responses towards several other aversive stimuli, including freezing, withdrawal, escape and avoidance.

VPMpc neurons have been known to send ascending projections to the IC. A few early studies using anterograde and retrograde WGA-HRP methods observed connections from VPMpc to rostrodorsal part of the LA in cats and rats^51–53^. We observed that VPMpc project their axon terminals to medial to posterior IC and rLA. With slice electrophysiology, we also verified the functional connection between VPMpc and these two brain regions, with the response of IC neurons slightly stronger than that of the rLA neurons. The axonal projection of VPMpc neurons to the IC layers was prominent in layer 1 and layer 3-4. The synaptic properties and organization of VPMpc afferents in different IC layers has been investigated by others; layer 4 neurons had maximal light-evoked EPSCs from VPMpc but some neurons in all layers responed^54^. While the IC has a well-established role in gustatory signal processing, whether individual insular neurons respond exclusively to one tastant or multiple tastants is controversial^55–57^. Non-gustatory functions of VPMpc projections to IC have not been studied to our knowledge. The VPMpc and IC have reciprocal connections; it also receives inhibitory input from the reticular thalamic nucleus^11,54^. Application of contemporary retrograde tracing techniques may reveal more inputs to molecularly defined VPMpc neurons. The LA is important in fear memory formation and consolidation; synaptic plasticity in LA is necessary to associate the conditional stimulus and the unconditional stimulus during fear memory formation^58,59^.

Thus, investigating how the projection from VPMpc to LA contributes to fear learning deserves examination. Together, our data expand the role of the VPMpc from a gustatory brain region to one that mediates a range of aversive behaviors.

## METHODS

### Animals

All mice used in this study were backcrossed onto a C57BL/6J background for greater than 6 generations. Homozygous *Cck^Cre/Cre^* mice were obtained from Jackson Laboratory (#012706). Homozygous *Oprm1^Cre/Cre^*and *Calca^FLPo/FLPo^* mice were generated and maintained as previously described^19,24^. Calca^FLPo/+^::Cck^Cre/+^ and Calca^FLPo/+^:: Oprm1^Cre/+^ mice were generated by breeding Calca^FLPo/FLPo^ with Cck^Cre/Cre^ or Oprm1^Cre/Cre^ mice, respectively. Heterozygous Oprm1^Cre/+^ were generated by breeding *Oprm1^Cre/Cre^* with wide-type C57BL/6J mice. All mice were maintained on a 12-h light/dark cycle (7 am-7 pm) with food and water *ad libitum* in a temperature- and humidity-controlled animal facility at the University of Washington. Mice were group-housed (3-5 mice per cage) and single-housed after stereotaxic surgery. All experiments were performed during the light cycle and mice were randomized to experimental groups with both male and female mice from the same litter. No sex differences were noted. All animal experimental protocols were approved by the Institutional Animals Care and Use Committee at the University of Washington (Protocol #2183–02).

### Virus production

Plasmids for pAAV1-Ef1ɑ-DIO-mCherry and pAAV1-hSyn-DIO-hM3Dq:mCherry were provided by B. Roth (Addgene #50462, #44361), pAAV1-Ef1ɑ-DIO-YFP and pAAV1-Ef1a-DIO-ChR2:YFP were provided by K. Deisseroth (Addgene #27056, #35507). pAAV1-Ef1ɑ-DIO- Synaptophysin:mCherry DNA plasmid was generated from pAAV1-Ef1ɑ-DIO-Synaptophysin:GFP with replacement of GFP^60^. AAV_DJ_-Ef1a-DIO-GFP:TetTox, AAV1-Ef1a-DIO-GCaMP6m and the FLP recombinase-dependent AAV1-CBA-frtDIO-ChR2-YFP were generated by R. Palmiter. All viruses except for AAV_DJ_-Ef1a-DIO-GFP:TetTox were prepared in-house by transfecting HEK cells with plasmids and pDG1 helper plasmids to coat AAV1 stereotype viruses. Viruses were then purified by mixing with sucrose and by CsCl-gradient ultracentrifugation. Viral pellets were re- suspended in 0.1 M phosphate-buffered saline (PBS) to yield approximately 10^13^ viral particles per mL. AAV_DJ_-Ef1a-DIO-GFP:TetTox was prepared by the Janelia Viral Tools lab.

### Stereotaxic surgery

Mice were anesthetized with isoflurane (5% for induction, 2% for maintenance, flow rate 1L/min) and placed on a robotic stereotaxic frame (Neurostar, Germany) for the entire surgery procedure. Mice also received local anesthesia of 1-2 mg/kg lidocaine and bupivacaine and skulls were surgically exposed. For calcium imaging recordings *in vivo*, a craniotomy was made unilaterally to target the VPMpc. AAV1-Ef1a-DIO-GCaMP6m virus (0.3 μl) was injected at a rate of 0.1 mm/min into the VPMpc (AP: -1.9 mm, ML: ± 0.7 mm, DV: 4.1 mm) and a 0.6 x 7.3 mm Integrated ProView GRIN lens (Inscopix, #1050-004413, California, USA) was placed above the virus injection target. For *in vivo* optogenetic behavior recordings, 0.3 μl AAV1-Ef1a-DIO- ChR2:YFP virus was unilaterally injected into the VPMpc with the same coordinates and an optic ferrule was placed 300 μm above the virus injection sites. Super glue (Loctite, Ohio, USA), dental cement (Lang Dental, Illinois, USA) and C&B metabond (Parkell, New York, USA) were used to secure both GRIN lens and optic ferrules. Other virus injection experiments were either targeted to the VPMpc with the same coordinates or PBN (AP: -4.9 mm, ML: ± 1.35 mm, DV: 3.4 mm) with a volume of 0.3 μl to VPMpc and 0.5 μl to PBN at a rate of 0.1 μl/min for 2 min using a Hamilton syringe (Reno, Nevada, USA). For nanoject viral injection surgeries, mice were placed on a stereotaxic frame (David Kopf Instruments, California, USA) and a 15 nl viral mix of AAV1-Ef1a-DIO-ChR2:YFP and AAV1-Ef1ɑ-DIO-Synaptophysin:mCherry was unilaterally injected into VPMpc with the same coordinates. We waited 10 min before withdrawing the injection needle. Following surgery procedures, 5 mg/kg Ketoprofen for analgesia was subcutaneously injected into mice which were monitored until complete recovery from anesthesia. All mice had a minimum 4 weeks of recovery before the start of behavioral experiments. Histological verification of viral injections and GRIN lens/ fiber optic placements were performed at the end of each experiment.

### Electrophysiology

#### Slice preparation

All mice were anesthetized with Euthasol (1 ml per 10 g body weight, i.p.) and intracardially perfused with 95% O_2_/5% CO_2_ saturated ice-cold cutting solution containing (in mM): 92 N-methyl-D-glucamine, 25 D-glucose, 2.5 KCl, 10 MgSO_4_, 1.25 NaH_2_PO_4_, 30 NaHCO_3_, 0.5 CaCl_2,_ 20 HEPES, 2 thiourea, 5 Na-ascorbate, 3 Na-pyruvate. After the intracardial perfusion, the mice were decapitated and the brain was removed and stored in the same ice-cold cutting solution superfused with 95% O_2_/5% CO_2_ saturated ACSF. Coronal slices (300 μm) were prepared using a vibratome (Leica VT1200, Illinois, USA) in the same ice-cold cutting solution. Brain slices were collected into the 33°C cutting solution for 10 min and then transferred to a room temperature recovery solution containing (in mM): 13 D-glucose, 124 NaCl, 2.5 KCl, 2 MgSO_4_, 1.25 NaH_2_PO_4_, 24 NaHCO_3_, 2 CaCl_2_, 5 HEPES for at least 1 hour. Brain slices were then individually transferred to a 33°C artificial cerebral spinal fluid (ACSF) containing (in mM): 11 D-glucose, 126 NaCl, 2.5 KCl, 1.2 NaH_2_PO_4_, 26 NaHCO_3_, 2.4 CaCl_2_, 1.2 MgCl_2_ for whole-cell patch recordings. All cutting, recovery, and ACSF solutions were saturated with 95% O_2_/5% CO_2_ and adjusted to pH 7.3-7.4, 300-310 mOsm.

#### Data acquisition and analysis

All recordings were obtained with glass pipettes (3-5 MΩ) filled with either Cs^+^ or K^+^ intracellular solutions. Cell-attached measurements were used to validate the efficiency of CNO applications using K^+^ intracellular solutions containing (in mM): 135 K- gluconate, 4 KCl, 10 HEPES, 4 Mg-ATP, 0.3 NA-GTP (pH 7.35, 280 -300 mOsm). Action potentials were recorded in voltage clamp with 0-pA holding current before and after bath applied 3 μM CNO onto thalamus slices with hM3Dq expression. To verify the connectivity between innervated neurons, light-evoked post-synaptic currents were recorded with Cs+ intracellular solutions containing (in mM): 120 CsMeSO_3_, 20 HEPES, 0.4 EGTA, 2.8 NaCl, 2.5 Mg-ATP, 0.25 Na-GTP, 5 QX-314 bromide (pH 7.35, 280-300 mOsm). For PBN-VPMpc connectivity verification, whole-cell patch-champ was performed on mCherry-positive VPMpc neurons after identifying the YFP expression in the PBN cell body and seeing dense green fibers in VPMpc with epifluorescence microscopy (OLYMPUS BX51WI, Evident, Massachusetts, USA). For measuring VPMpc-IC and VPMpc-LA connectivity, only neurons with dense YFP fibers from VPMpc were patched. All neurons were recorded in voltage clamp with -70 mV holding potential and the post-synaptic currents were evoked by a 0.1-Hz 10-ms blue light pulsed delivered through the 40× objective via a SOLIS-3C high power LED (Thorlabs, New Jersey, USA) and filter cube (Chroma technology, Vermont, USA). Averaged data were obtained from 10 sweeps. All data were obtained using MultiClamp 700B amplifier (Molecular Devices, California, USA). Data acquisition and analysis were done using pClamp 11 and Clampfit 11.3 (Molecular Devices, California, USA).

### Optogenetic stimulation

After allowing 6 weeks to recover from viral surgery and optic ferrule implantation, dummy cables were attached to acclimate mice prior to testing. For behavioral studies, mono 200-μm diameter fiber-optic cables (Doric Lenses, Québec, Canada) were attached to the optic ferrule of each mouse before experiments. A blue-light laser (473-nm; LaserGlow, Ontario, Canada) was used to stimulate ChR2 (10-ms light-pulse trains were delivered at 10 Hz, 2 ± 0.5 mW) using a Master-8 pulse stimulator (AMPI, Maryland, USA) in all behavioral tests. For von Frey-testing using optogenetic stimulation, 5 min of light stimulation (as above) was delivered in home cages 45 min before the tactile stimulation tests.

### Calcium Imaging

Mice were prepared and recorded as previously described^43,61^. After allowing 6 weeks to recover from GCaMP6m virus injection and GRIN lens implantation surgery, each mouse’s head was attached to a micro endoscope (Inscopix, California, USA) and connected to nVista 3.0 (Inscopix, California, USA) to acquire calcium images. Ethovision XT15 (Noldus, Virginia, USA) and Med Associates control panel (Med Associates, Vermont, USA) were connected to nVista 3.0 via BNC cables to trigger TTL pulses to allow synchronizing calcium signals with behavioral video recordings and conditional and unconditional stimuli. All mice recorded were freely moving and without any anesthetization. A commutator (Inscopix, California, USA) was used to avoid the overspinning of the micro endoscope during recording.

For sensory stimuli, mice were placed in their home cage and went through multiple trials of a transient 90 dB hand clapping sound, air puff on the face, tail pinch, and 10 s of lifting the mouse by the tail 0.5 m within one recording session, with a random ITI of 60-120 s.

Heatmaps of sensory stimuli used the first trials for each mouse. For foot shock only recordings, mice were placed in the shock chamber and allowed 5 min free exploration before shock session started. During the shock session, mice went through 1-2 trials of 0.15, 0.3, 0.5 mA foot shock stimulation with an averaged ITI of 120s. For fear conditioning recordings, mice were placed in the test chamber 5 min before calcium imaging starts across all three days. On Day 1, 10 trials of 10 s, 5 kHz, 65 dB tone were delivered with a 100-s ITI. On Day 2, 10 trials of 10 s, 5 kHz, 65 dB tone were co-terminated with 0.5-s, 0.5-mA foot shocks (ITI 100 s). On Day 3, mice were placed in a different chamber and only 10 trials of 10 s, 5 kHz, 65 dB tone were delivered with an ITI of 100 s.

Calcium fluorescence was recorded at 20 frames per second and under LED light while mice were being tested. The recording parameters were chosen from pilot recordings with minimal photo-bleaching but sufficient detection of fluorescence. Raw calcium recording videos were pre-processed with Inscopix Data Processing Software (IDPS 1.9.1, Inscopix, California, USA) with 2× down sampling of frame and spatial bandpass filter to reduce processing time and background noise. Rigid motion correction algorithm provided by IDPS was also applied to the filtered images to minimize motion artifact during analysis. Constrained non-negative matrix factorization for microendoscope data (CNMF-E) analysis was used to extract the raw F over noise of each neuron within the region of interest in field of view^62^. All CNMF-E detected calcium transients were visually inspected for each cell to ensure accuracy. Raw traces (F over noise, equivalent to F) after CNMF-E were analyzed using custom code in Matlab to generate ΔF/F and Z score. ΔF/F was calculated as ΔF/F=([F-mean(F0)])/(mean(F0)). Z score was calculated as Z=([F-mean(F0)])/(Standard deviation(F0)). F indicates the fluorescence at any given point and F0 indicates the average fluorescence of -20 to 0 second in fear conditioning recordings and -10 to 0 second in all other recordings.

The area under the curve (AUC) of ΔF/F were used to identify whether the cell was activated, inhibited or unresponsive. AUC was calculated for both pre-stimulation (-10 s to 0 s) and post-stimulation (0 s to 10 s) across all trials. Neurons were deemed as ‘increased’ if the AUC of post-stimulation has > 50% increase compared to the AUC of pre-stimulation. Neurons were deemed as ‘decreased’ if < 50% and redeemed as “no response” if the difference of AUC did not show > 50% increase and decrease.

### Behavior measurements

#### Open-field test

Mice injected with YFP or ChR2 were attached to fiber-optic patch cords and placed in a 40 cm × 40 cm × 30 cm white plexiglass open field box for at least five minutes of habituation. Photostimulation was for 30 s (473 nm, 10 Hz and 2 mW) and delivered 3 times with one-minute intervals between stimulations. Mice were attached with fiber-optic patch cords for both control and experimental groups to minimize the variation. The distance traveled was collected and analyzed using video-tracking software EthoVision XT 15 (Noldus, Virginia, USA).

#### Von Frey test

The von Frey tactile sensitivity test was performed as described^43^. Mice were placed in a 30-cm elevated metal mesh floor with 11.5 cm×7.5 cm grid separated by black plexiglass (Bioseb, Florida, USA) for at least 30 min before experiments. Von Frey filaments (Bioseb, Florida, USA) with different stiffness ranging from 0.07 to 4 g were used to measure the tactile sensitivity of the hindpaw plantar surface in each mouse. Filaments were applied from the minimum weight and switched to a higher weight if there were less than 3 paw- withdraw responses within 5 trials. The lowest weight that elicited 3 paw-withdraw responses with 5 trials was recorded as paw withdraw threshold and confirmed with the next consecutive filament. The ipsilateral and contralateral paw-withdrawal thresholds were separately measured and combined in the data analysis due to no differences observed in left/right paw-withdrawal thresholds.

#### Hot-plate test

*A* hot-plate test was conducted as described^43^. Mice were gently placed on a 52.5 ± 1 °C pre-heated 16.5 × 16.5 cm aluminum surface embedded in a 16.5 × 16.5 × 35 cm rectangular clear plexiglass enclosure (Bioseb, Florida, USA). The latency to the first jump and the number of jumps were recorded and measured in 60-s trial tests. Additional time (maximum 30 s) were given to mice resisted to jump in the 60-s trial until it jumped. For individuals resisted to jump, the animal was immediately removed from the hot plate after 90 s to avoid tissue damage.

#### Fear conditioning

Mice were placed in a fear-conditioning chamber consisting of a 20×14×13 cm shuttle box with a metal grid floor, a light on the wall and a speaker in the back (Med Associates, Vermont, USA). The metal grid floor is made of stainless-steel bars and delivers an electric shock. The metal grid floor, light and speaker were all connected and controlled by Med Associates software. A 3-day series of trials were conducted. Day 1: Mice were placed in the chamber and allowed free exploration for 60 s before the onset of a discrete conditioned stimulus, which consisted of 6 trials with a 20 s, 5 kHz, 65 dB tone with a 100-s inter-trial interval (ITI). Day 2: Mice were allowed to explore the apparatus for 60 s and 6 trials of 20 s, 5 kHz, 65 dB CS tone was delivered that co-terminated by a 2-s, 0.5-mA foot shock with an ITI of 100 s. After the sixth CS-US paring trial, mice remained in the chamber for another 100 s before returning to the home cage. Day 3: For the contextual test, mice were placed in the same chambers for 2 min of free exploration. For the cue test, mice were placed in a different chamber and allowed to explore the cage for 60 s and then one CS tone (20 s, 5 kHz, 65 dB) was delivered. Freezing data were recorded in a camera synchronized with the Med Associates software and freezing time (immobility time with both mouse head and body) during each trial was manually scored by the examiner.

#### Fear-potentiated startle

This test was performed as described^63^. Mice were placed in the standardized enclosure of a sound-attenuating startle reflex chamber (SR-LAB, San Diego Instruments, California, USA) for 5 min of free exploration with a background of 65 dB white noise. 7 trials of escalating sound from 65 (null), 80, 90, 100, 105, 110 and 120 dB were presented to mice with an ITI of 30 s. This 7-trial series was repeated 10 times with the same ITI. All sounds were delivered with 40-ms pulses. Startle responses upon each sound level changer were calculated and averaged for all 10 repeated trials. For fear-potentiated startle, mice undergo a 3-day protocol. On day 1 (baseline day), mice were pseudo-randomly given 10 trials of cue and 10 trials of no-cue conditions after a 5-min habituation period. For cue trials, mice were presented with 10-s light and a 40-ms, 105-dB sound pulse co-terminating with the light. For no-cue trails, mice were only presented with 40-ms, 105-dB sound pulse without any light cue exposure. The ITI was 60 to 180 s with an average of 120 s. On day 2 (training day), after a 10-min habituation period in the startle chamber, mice were given 30 trials of 10-s light cue stimulation, co-terminated with 0.2 mA, 0.5 s foot shock. The ITI between these 30 trials also ranged from 60 to 180 s, with an average of 120 s. On day 3 (test day), mice underwent the same protocol as on Day 1. All no-cue and cue trial responses were averaged separately on Day 1 and Day 3. Fear-potentiated startle was calculated as [(averaged responses on cue trials / average responses on no-cue trials - 1) × 100].

#### Conditioned place preference (CPP)

Mice underwent a 4-day conditioned place preference paradigm protocol in a rectangular CPP box. The CPP box is divided equally into two 20×20×30 cm square boxes with a 6.5×30 cm slide door connected in the middle. These two square boxes differ in wall decoration with vertical stripes one side and spots on the other side. The vertical striping chamber also contains two triangular blocks in the corner. On day 1, Mice were placed in the middle of the CPP chamber and allowed free exploration for 30 min with open access to both striped and dotted chambers. On day 2 and day 3, mice were i.p. injected with Saline (10ml/kg) and immediately moved to the saline-paired chamber (either the vertical stripe or circled dot chamber) with the door closed. Mice were allowed free exploration in the saline-paired chamber for 30 min. Five hours later, mice received the same volume of U50 (5 mg/kg) through i.p injection and were moved immediately to the other chamber paired with U50. Mice were allowed free exploration in the U50-paired chamber for 30 min and returned to their home cage after. On day 4, mice were placed in the middle of two chambers and allowed to freely explore both sides for 10 min. The time spent on each side was measured and averaged using the video-tracking software, EthoVision XT 15 (Noldus, Virginia, USA), to calculate the conditioned preference.

#### Real-time, place preference (RTPP)

RTPP test was performed as previously described^25^. The RTPP box is a rectangular box divided into two equal-sized square box (60×30×30 cm) with one side painted with black dots on the wall and the other side being blank without any symbols.

The dotted and blank chambers were connected through a single doorway (10×30 cm) in the middle. Mice injected with YFP or ChR2 were attached to fiber-optic patch cords and underwent two days of RTPP test paradigm. On day 1, mice were placed in the middle of the two chambers and allowed free exploration for 20 min without any light stimulation. On day 2, mice were also placed in the middle of the chamber and allowed to be freely moving for 20 min, but one side of the chamber was paired with optical stimulation. Mice received 473 nm 10 Hz 2 mW light stimulation once it moved to the stimulation chamber and the light stimulation was continuously turned on until it moved out of the light-paired chamber. The time spent on the light-paired chamber and the no-light chamber was measured and averaged using the video-tracking software, EthoVision XT 15 (Noldus, Virginia, USA), to calculate the real-time place preference.

#### RNAscope *in situ* hybridization

Mice injected with AAV expressing hM3Dq were given CNO (1 mg/kg, i.p.) and put back in their home cage. After 60 min, mice were anesthetized with Euthasol (1 ml per 10 g body weight) and decapitated. Brains were quickly removed and rapidly frozen on crushed Dry Ice. coronal cryostat sections (20 μm) were cut on cryostat (Leica Biosystems, Illinois, USA) and directly transferred to a SuperFrost Plus glass slide (Thermo Fisher Scientific, Massachusetts, USA) and frozen at -80°C. RNAscope fluorescent multiplex V1 assay was performed following the manufacturer’s protocols (ACD, California, USA) using the following probes: *Cck*, *Oprm1* and *Fos* (ACD, California, USA). All images were collected on Keyence BZ-710 microscope (Keyence, Illinois, USA) and Olympus FV-1200 confocal microscope (Evident, Massachusetts, USA). RNA expression and colocalization were analyzed using Fiji with manually drew regions of interest.

Cells with at least 4 puncta associated with a DAPI nucleus expression were considered positive.

### Histology and immunohistochemistry

The detailed procedures for mice brain slice preparation were described^20,43^. Mice were anesthetized using pentobarbital (0.2 ml, i.p.) and transcranial perfused with 1 × ice-cold phosphate-buffered saline (PBS) followed by 4% ice-cold paraformaldehyde (PFA, Electron Microscopy Sciences, Pennsylvania, USA) made in PBS. For virus injection verification, each brain was rapidly removed and fixed overnight in 4% PFA at 4°C. For optic ferrule and lens implantations, the whole mouse head was decapitated and moved to 4% PFA at 4°C for at least 24 h and the brain was then dissected and post-fixed in 4% PFA at 4°C for 6 h. Then the brain was transferred to 30% sucrose for 24-36 h until it was fully saturated and sank. The brain was frozen in OCT compound (Thermo Fisher Scientific, Massachusetts, USA) and stored at -80°C. Before sectioning, the brain was transferred from -80°C to -20°C for at least 30 min to prepare for the cryostat sectioning (Leica Biosystems, Illinois, USA). Coronal cryostat sections (35 μm) were cut on cryostat and collected in ice-cold PBS.

For immunohistochemistry, sections were washed 3 times in 1 × PBS and blocked with 3% normal donkey serum made in PBST (0.2% Triton X-100 in PBS) for 1 h at room temperature. Sections were then incubated overnight at 4°C in a blocking solution with primary antibodies including chicken-anti-GFP (1:10000, Abcam, ab 13970) or rabbit-anti-dsRed (1:2000, Takara, ab 632496). After washing the residual primary antibodies with PBS for 3 min the following day, sections were incubated at room temperature for an hour in PBS with secondary antibodies including Alexa Fluor 488 donkey anti-chicken (1:500, Jackson ImmunoResearch) and Alexa Fluor 594 donkey anti-rabbit (1:500, Jackson ImmunoResearch). Following washing with PBS for 3 times, the sections were mounted onto SuperFrost Plus glass slides (Thermo Fisher Scientific, Massachusetts, USA) and coverslipped with Fluoromount-G with DAPI (Southern Biotech, Alabama, USA). Fluorescent images were collected on Keyence BZ-710 microscope (Keyence, Illinois, USA) and Olympus FV-1200 confocal microscope (Evident, Massachusetts, USA). Images were minimally processed using Fiji to enhance brightness and contrast. All images were processed the same way.

### Statistical Analyses

All the averaged data are shown as mean ± standard error of the mean (SEM). All data were evaluated by independent *t*-tests, paired *t*-tests, two-way repeated measures ANOVA, followed by Post-hoc Holm-Sidak’s multiple comparisons when appropriate. P < 0.05 was considered as statistically significant and corresponding to the following values: *P < 0.05, **P < 0.01, ***P < 0.001. Independent sample size (n) represented the number of animals in behavioral tests, the number of brain slices in RNAscope analysis, and number of neurons in electrophysiological recordings. Mice were excluded from experimental analysis if the viral expression was inadequate, or off-target based on histology and imaging verification. All analyses were conducted using GraphPad PRISM (GraphPad Software, California, USA) and MATLAB (MathWorks, Massachusetts, USA)

## ACKNOWLEDGMENTS

We thank Susan Phelps and Lucy Anastas for maintaining the mouse colony, Larry Zweifel and his lab members for insightful discussions, An-Doan Nguyen for her assistance in data analysis, Anqi Zhu for technical assistance in MATLAB analysis and all members in Palmiter lab for their comments on the manuscript.

## AUTHOR CONTRIBUTIONS

F.C. and R.D.P. conceived and designed the study. F.C. and S.P. performed and analyzed the 1- photo calcium imaging. F.C performed and analyzed the electrophysiology experiments. F.C. and E.Y.S. performed and analyzed the behavioral experiments. F.C, J.L.P, E.Y.S performed and analyzed the histology experiments. F.C and R.D.P wrote the manuscript with input from all authors.

## DATA AVAILABILITY

Source data are provided with this manuscript. Other Data are available from the corresponding author upon reasonable request.

---

**Extended Data Fig. 1.**
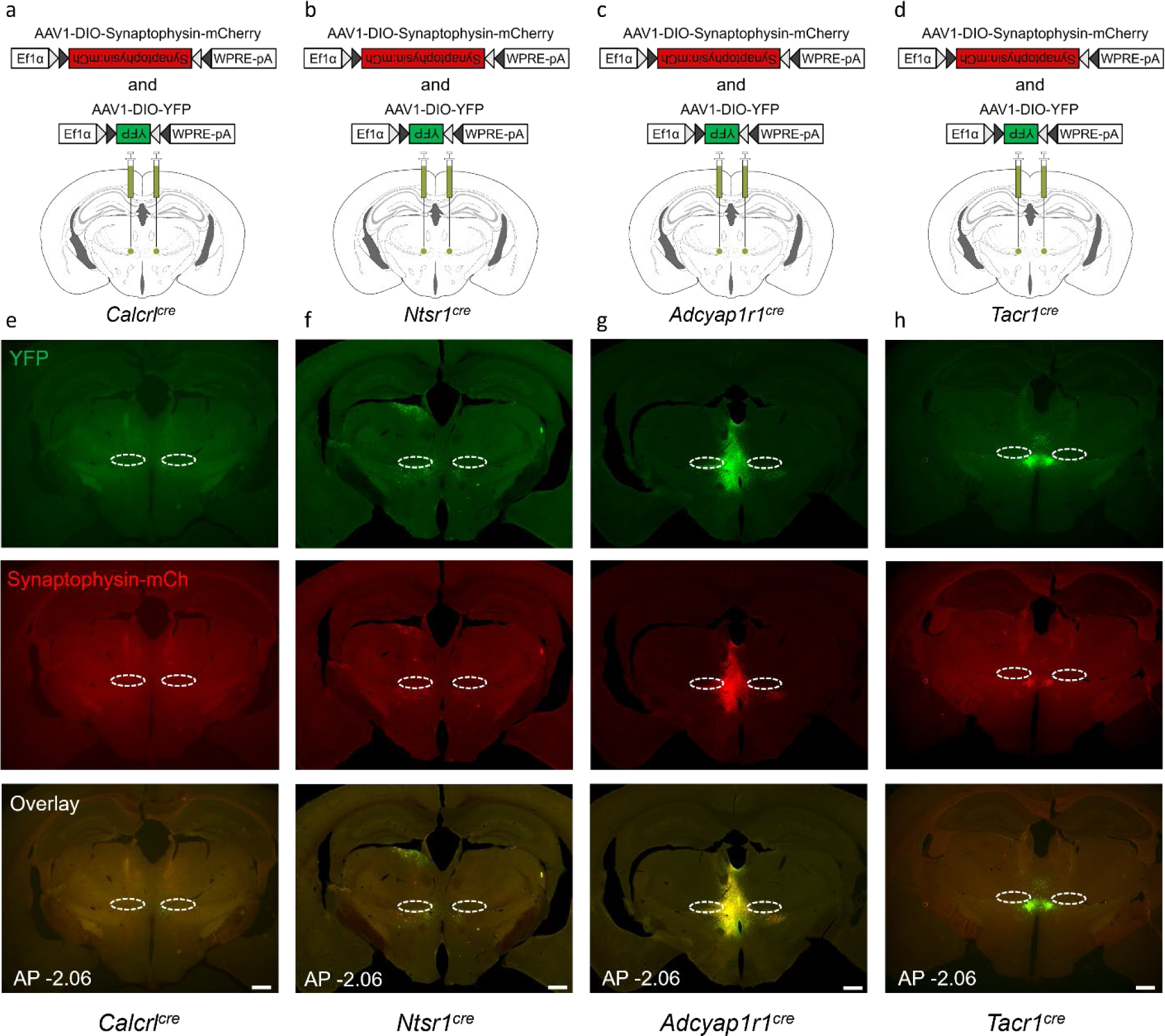
YFP and Synaptophysin expression in VPMpc of *Calcrl^Cre^, Ntsr1^Cre^, Crhr1^cre^, Adcyap1r1^Cre^* or *Tacr1^Cre^* mice. a-d. , Scheme showing bilateral injection of AAV1-DIO-mYFP and AAV1-DIO- Synaptophysin:mCherry into the VPMpc of *Calcrl^cre^* (**a**), *Ntsr1l^cre^* (**b**), *Adcyap1r1^cre^* (**c**), and *Tacr1l^cre^* (**d**) mice. **e**, YFP (top), Synaptophysin:mCherry (middle) expression in the VPMpc of *Calcrl^cre^* mice. **f**, YFP (top), Synaptophysin:mCherry (middle) expression in the VPMpc of *Ntsr1l^cre^*mice. **g**, YFP (top), Synaptophysin:mCherry (middle) expression in the VPMpc of *Adcyap1r1^cre^* mice. **h**, YFP (top), Synaptophysin:mCherry (middle) expression in the VPMpc of *Tacr1l^cre^*mice. Scale bar 500 μm. AP: anterior-posterior bregma level, dotted circle is the VPMpc.

**Extended Data Fig. 2.**
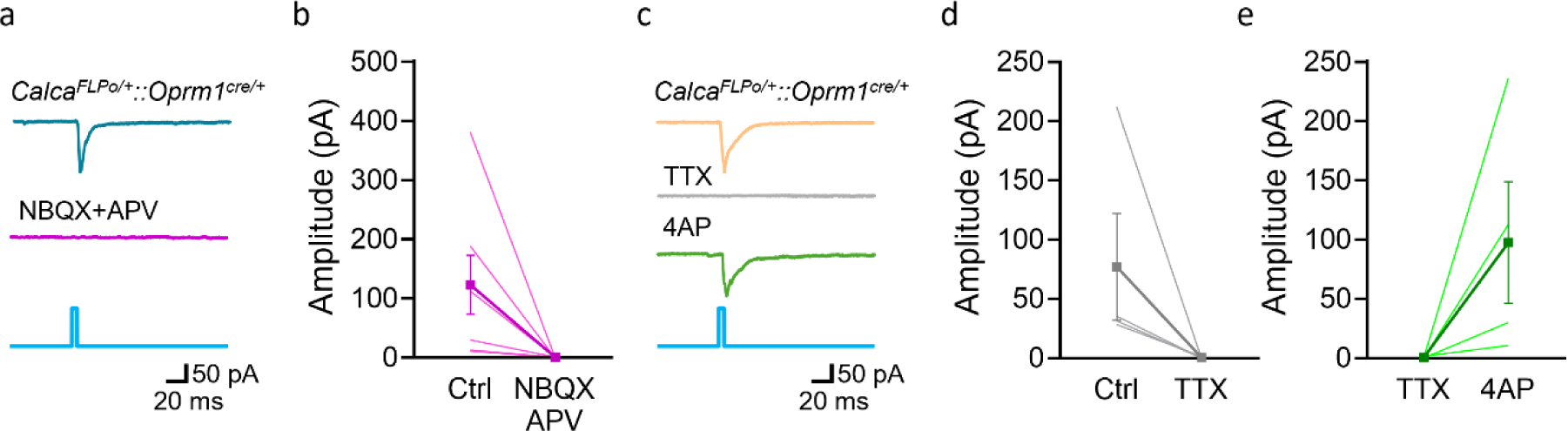
**CGRP^PBN^ neurons send direct excitatory input to OPRM1^VPMpc^ neurons a**, Sample traces and **b**, summary figure of EPSCs in the VPMpc neurons of *Calca^FLPo^::Oprm1^Cre^* evoked by 455-nm blue light illumination and blocked by NBQX and APV. **c**, Sample traces and summary figures of EPSCs in the VPMpc neurons of *Calca^FLPo^::Oprm1^Cre^* evoked by 455-nm blue light illumination, **d**, blocked by TTX and **e**, reinstated by 4AP. Data are represented as mean ± SEM.

**Extended Data Fig. 3.**
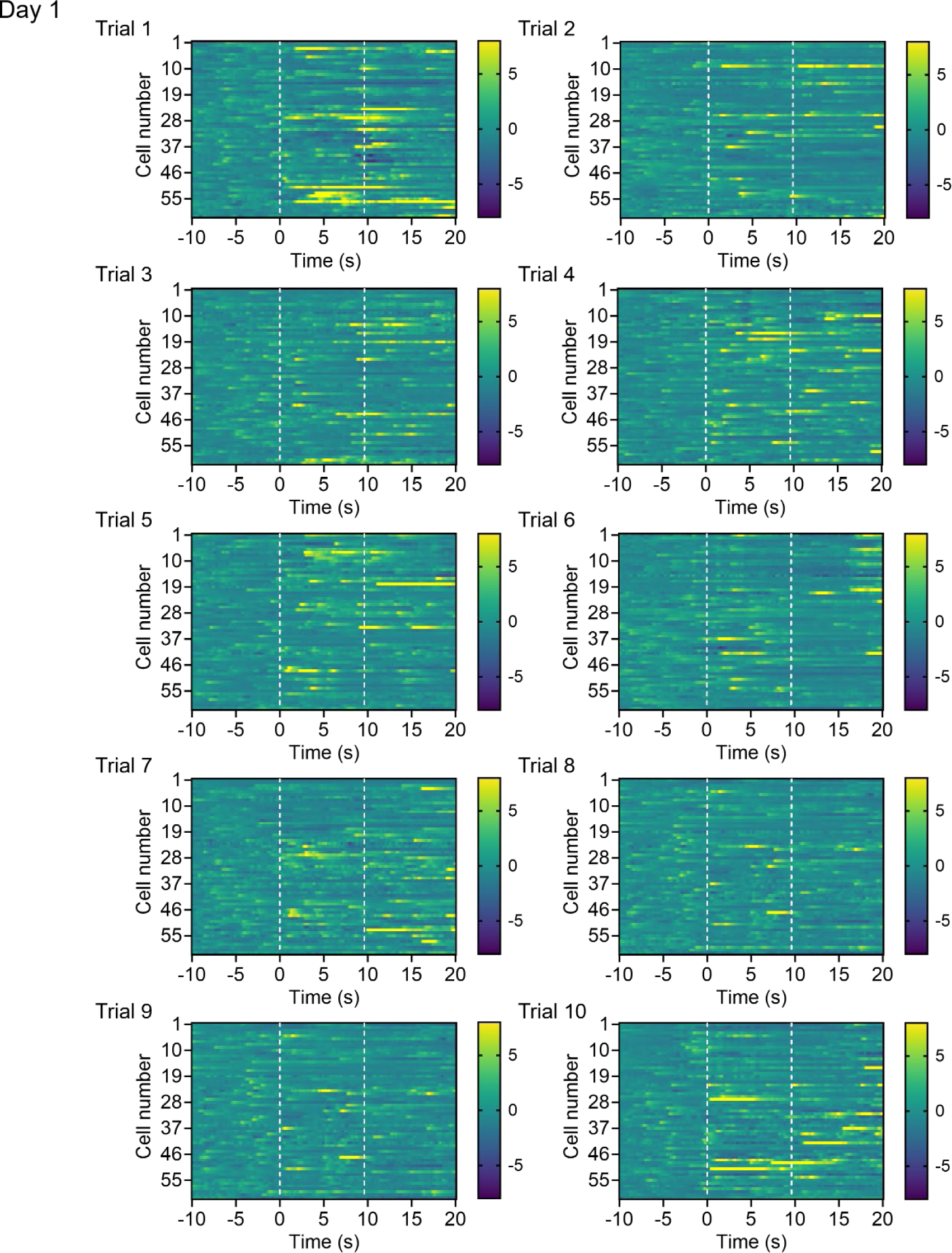
**Heat maps of calcium fluorescence activity during 10 trials of 30 s calcium recordings in Day 1**

**Extended Data Fig. 4.**
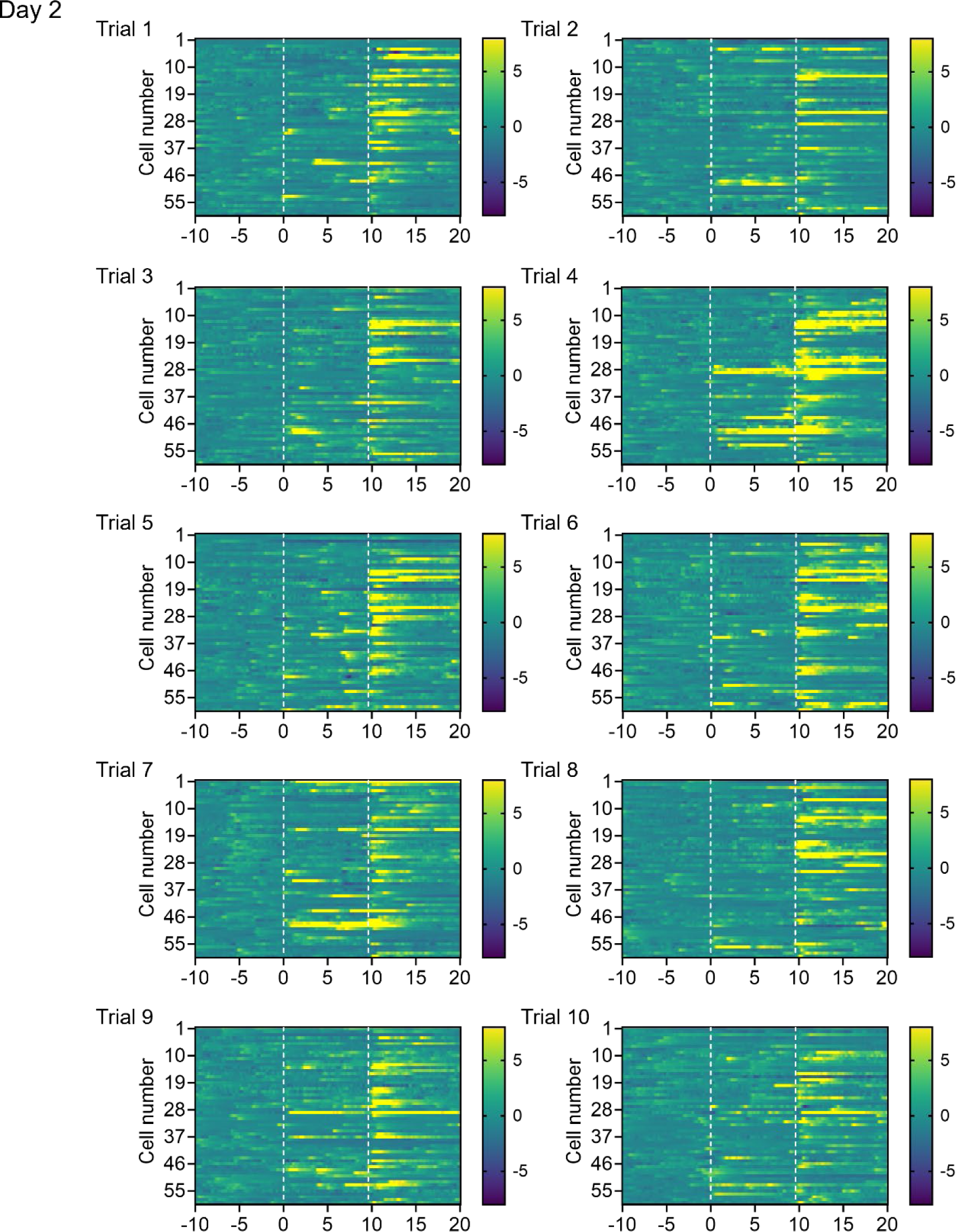
**Heat maps of calcium fluorescence activity during 10 trials of 30 s calcium recordings in Day 2**

**Extended Data Fig. 5.**
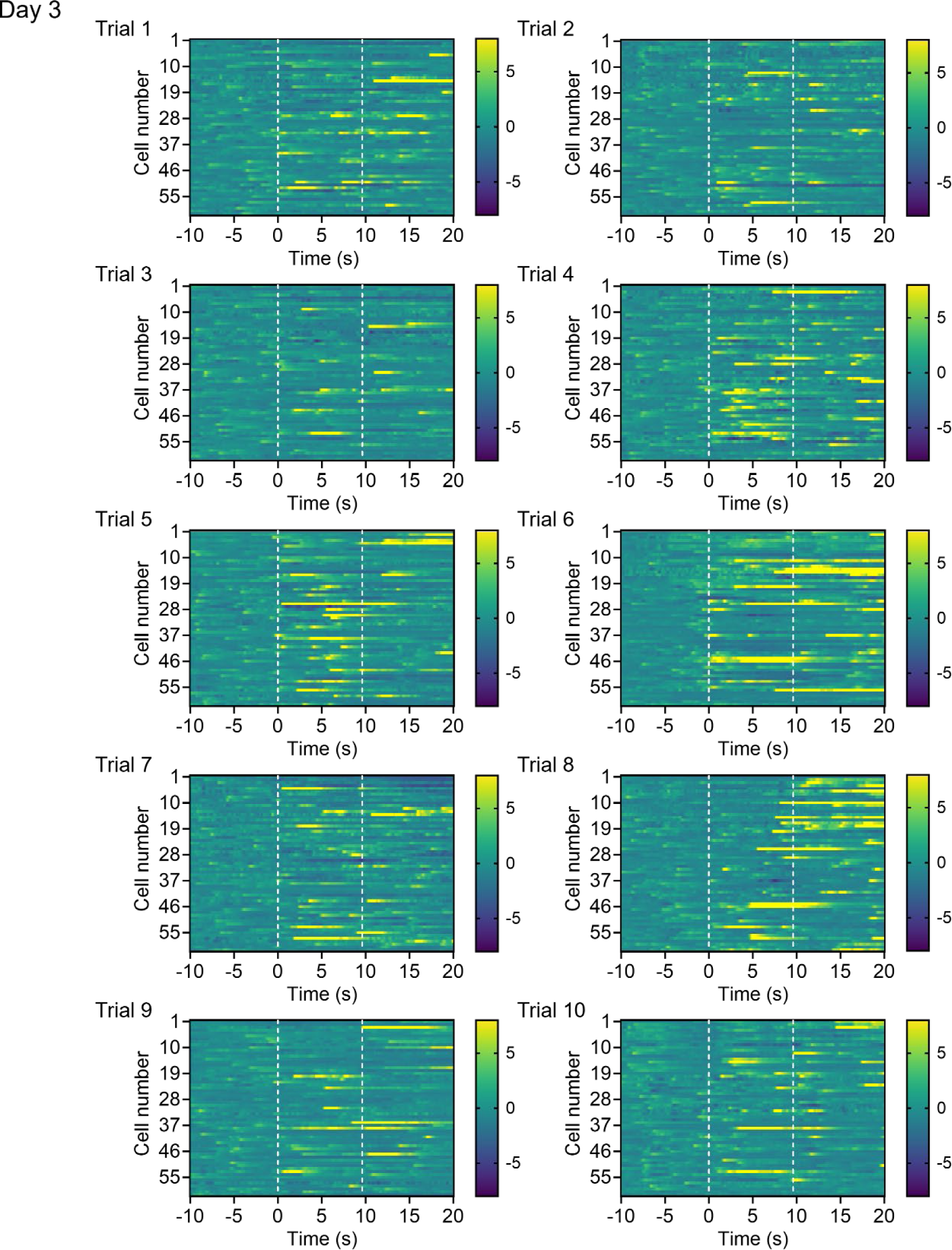
**Heat maps of calcium fluorescence activity during 10 trials of 30 s calcium recordings in Day 3**

**Extended Data Fig. 6.**
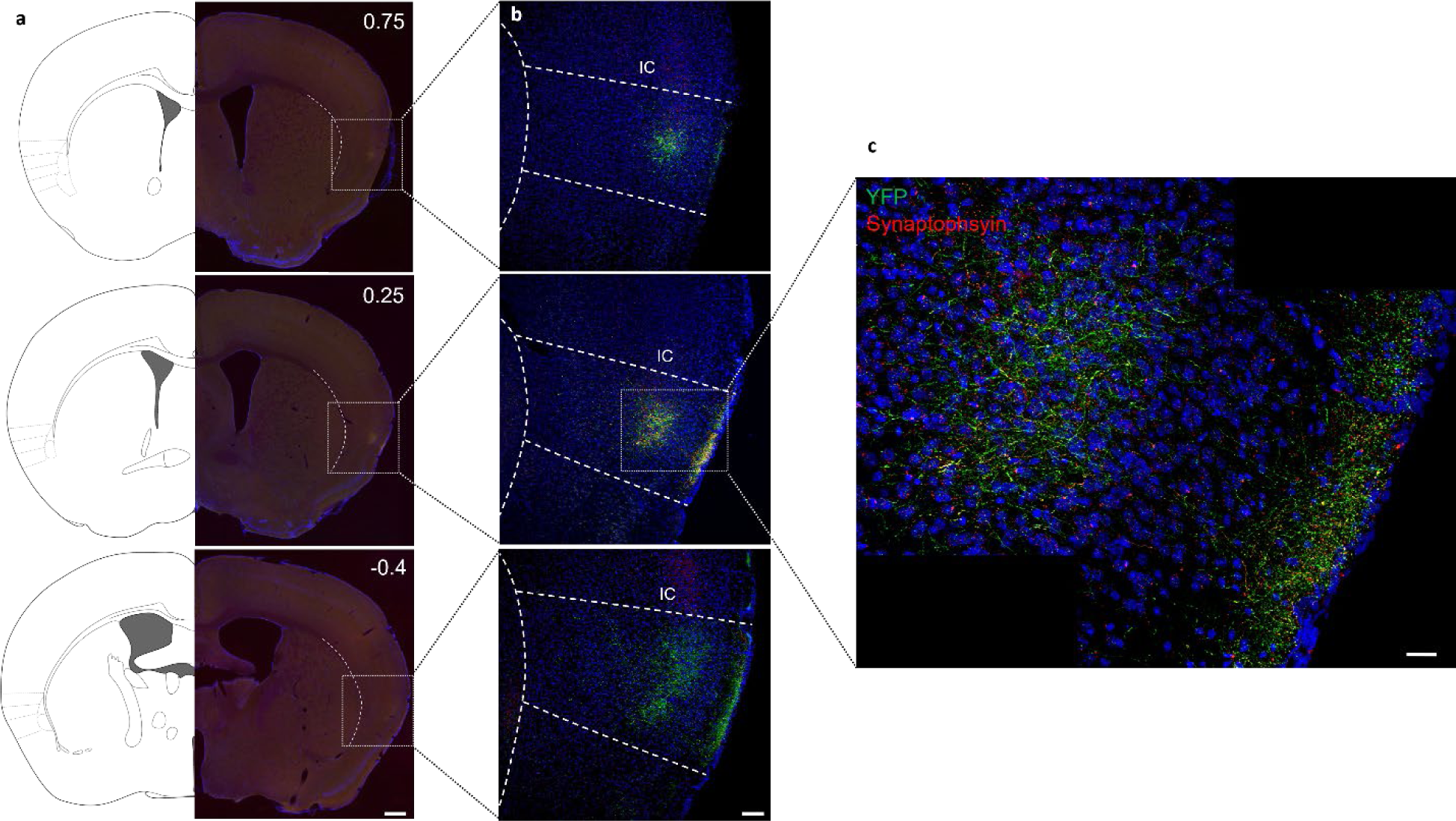
**CCK^VPMpc^ neurons send axonal projection to IC a**, Coronal map (left) and slice Images (right) showing VPMpc axon terminals in different bregma levels containing the IC, scale bar 500 μm. **b**, Higher magnification images of the white box in (**a**) showing VPMpc axon terminals in different bregma levels of IC, scale bar 100 μm. **c**, Zoom-in images of the white box in (**b**) showing VPMpc axon terminals expressed in IC (bregma 0.25), scale bar 25 μm.

**Extended Data Fig. 7.**
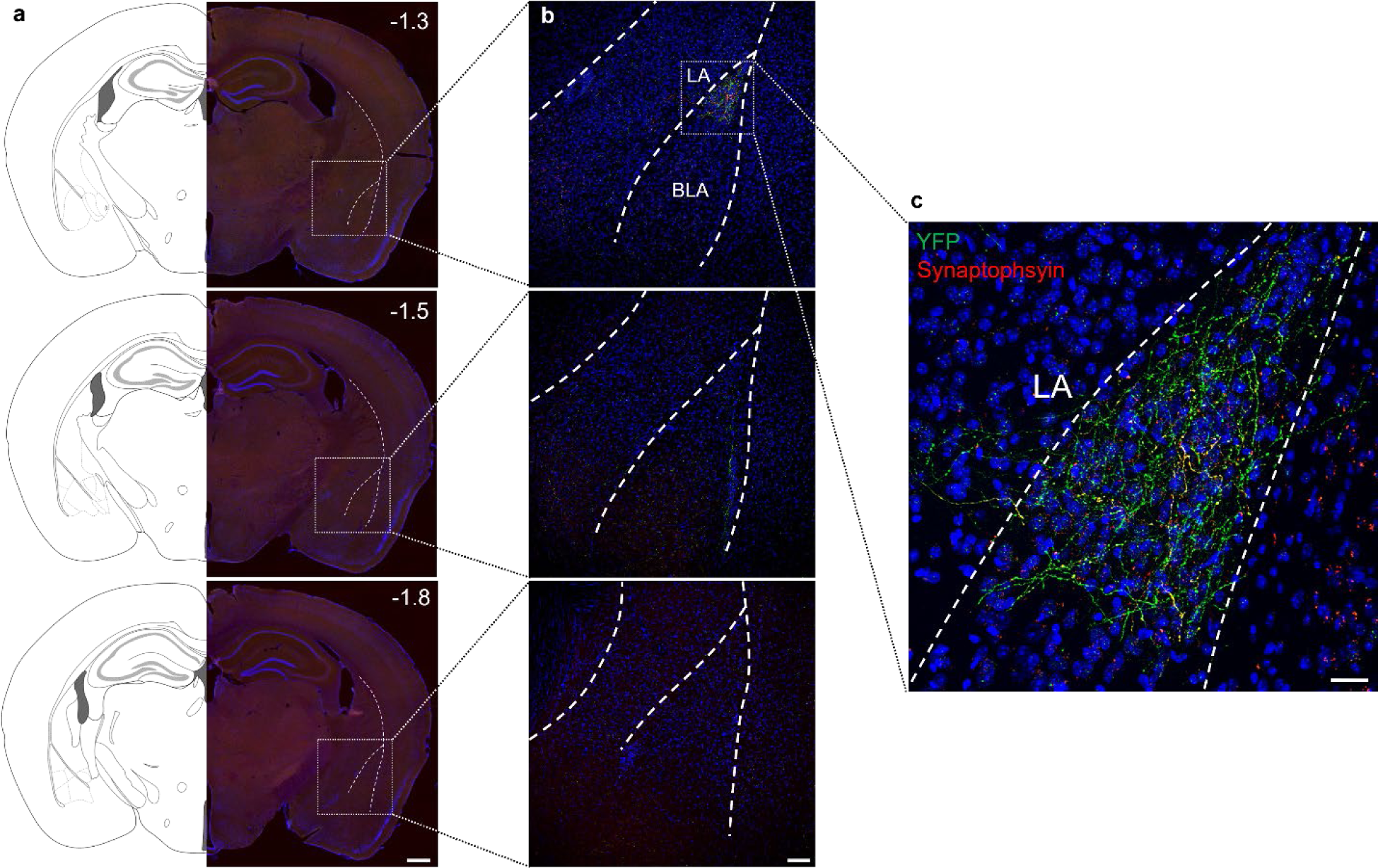
CCK^VPMpc^ neurons send axonal projection to rostral LA **a**, Coronal map (left) and slice Images (right) showing VPMpc axon terminals in different bregma levels containing the LA, scale bar 500 μm. **b**, Higher magnification images of the white dash box in (**a**) showing VPMpc axon terminals in different bregma levels of LA, scale bar 100 μm. **c**, Zoom-in images of the white dash box in (**b**) showing VPMpc axon terminals only expressed in rostral LA (bregma -1.3), scale bar 25 μm.

## REFERENCES

1. 1 Jones, E. The thalamus Plenum Press New York. (1985).

2 Purves, D. et al. Neuroscience 2nd edition. sunderland (ma) sinauer associates. Types of Eye Movements and Their Functions (2001).

3 Torrico, T. J. & Munakomi, S. Neuroanatomy, thalamus. (2019).

4. Jones, E. The Thalamus. Cambridge Univ Press. Cambridge, UK (2007).

5 Roy, D. S., Zhang, Y., Halassa, M. M. & Feng, G. Thalamic subnetworks as units of function. Nat Neurosci 25, 140–153 (2022). 10.1038/s41593-021-00996-1

6 Pritchard, T. C., Hamilton, R. B. & Norgren, R. Neural coding of gustatory information in the thalamus of Macaca mulatta. J Neurophysiol 61, 1–14 (1989). 10.1152/jn.1989.61.1.1

7 Samuelsen, C. L., Gardner, M. P. & Fontanini, A. Thalamic contribution to cortical processing of taste and expectation. J Neurosci 33, 1815–1827 (2013). 10.1523/JNEUROSCI.4026-12.2013

8 Verhagen, J. V., Giza, B. K. & Scott, T. R. Responses to taste stimulation in the ventroposteromedial nucleus of the thalamus in rats. J Neurophysiol 89, 265–275 (2003). 10.1152/jn.00870.2001

9 Beckstead, R. M., Morse, J. R. & Norgren, R. The nucleus of the solitary tract in the monkey: projections to the thalamus and brain stem nuclei. J Comp Neurol 190, 259–282 (1980). 10.1002/cne.901900205

10 Bester, H., Bourgeais, L., Villanueva, L., Besson, J. M. & Bernard, J. F. Differential projections to the intralaminar and gustatory thalamus from the parabrachial area: a PHA-L study in the rat. J Comp Neurol 405, 421–449 (1999).

11 Holtz, S. L., Fu, A., Loflin, W., Corson, J. A. & Erisir, A. Morphology and connectivity of parabrachial and cortical inputs to gustatory thalamus in rats. J Comp Neurol 523, 139–161 (2015). 10.1002/cne.23673

12 Karimnamazi, H. & Travers, J. B. Differential projections from gustatory responsive regions of the parabrachial nucleus to the medulla and forebrain. Brain Res 813, 283–302 (1998). 10.1016/s0006-8993(98)00951-2

13 Norgren, R. & Leonard, C. M. Ascending central gustatory pathways. J Comp Neurol 150, 217–237 (1973). 10.1002/cne.901500208

14 Katz, D. B., Simon, S. A. & Nicolelis, M. A. Dynamic and multimodal responses of gustatory cortical neurons in awake rats. J Neurosci 21, 4478–4489 (2001). 10.1523/JNEUROSCI.21-12-04478.2001

15 Samuelsen, C. L., Gardner, M. P. & Fontanini, A. Effects of cue-triggered expectation on cortical processing of taste. Neuron 74, 410–422 (2012). 10.1016/j.neuron.2012.02.031

16 Nomura, T. & Ogawa, H. The taste and mechanical response properties of neurons in the parvicellular part of the thalamic posteromedial ventral nucleus of the rat. Neurosci Res 3, 91–105 (1985). 10.1016/0168-0102(85)90024-0

17 Reilly, S. & Pritchard, T. C. Gustatory thalamus lesions in the rat: II. Aversive and appetitive taste conditioning. Behav Neurosci 110, 746–759 (1996).

18 Liu, H. & Fontanini, A. State Dependency of Chemosensory Coding in the Gustatory Thalamus (VPMpc) of Alert Rats. J Neurosci 35, 15479–15491 (2015). 10.1523/JNEUROSCI.0839-15.2015

19 Chen, J. Y., Campos, C. A., Jarvie, B. C. & Palmiter, R. D. Parabrachial CGRP Neurons Establish and Sustain Aversive Taste Memories. Neuron 100, 891–899 e895 (2018). 10.1016/j.neuron.2018.09.032

20 Pauli, J. L. et al. Molecular and anatomical characterization of parabrachial neurons and their axonal projections. Elife 11 (2022). 10.7554/eLife.81868

21 Fu, O. et al. SatB2-Expressing Neurons in the Parabrachial Nucleus Encode Sweet Taste. Cell Rep 27, 1650–1656 e1654 (2019). 10.1016/j.celrep.2019.04.040

22 Campos, C. A., Bowen, A. J., Roman, C. W. & Palmiter, R. D. Encoding of danger by parabrachial CGRP neurons. Nature 555, 617–622 (2018). 10.1038/nature25511

23 Han, S., Soleiman, M. T., Soden, M. E., Zweifel, L. S. & Palmiter, R. D. Elucidating an Affective Pain Circuit that Creates a Threat Memory. Cell 162, 363–374 (2015). 10.1016/j.cell.2015.05.057

24 Palmiter, R. D. The Parabrachial Nucleus: CGRP Neurons Function as a General Alarm. Trends Neurosci 41, 280–293 (2018). 10.1016/j.tins.2018.03.007

25 Bowen, A. J. et al. Dissociable control of unconditioned responses and associative fear learning by parabrachial CGRP neurons. Elife 9 (2020). 10.7554/eLife.59799

26 Gehrlach, D. A. et al. Aversive state processing in the posterior insular cortex. Nat Neurosci 22, 1424–1437 (2019). 10.1038/s41593-019-0469-1

27 Lin, J. Y., Arthurs, J. & Reilly, S. Gustatory insular cortex, aversive taste memory and taste neophobia. Neurobiol Learn Mem 119, 77–84 (2015). 10.1016/j.nlm.2015.01.005

28 Zheng, J. et al. An insular cortical circuit required for itch sensation and aversion. Curr Biol 34, 1453–1468 e1456 (2024). 10.1016/j.cub.2024.02.060

29 Lein, E. S. et al. Genome-wide atlas of gene expression in the adult mouse brain. Nature 445, 168–176 (2007). 10.1038/nature05453

30 Armbruster, B. N., Li, X., Pausch, M. H., Herlitze, S. & Roth, B. L. Evolving the lock to fit the key to create a family of G protein-coupled receptors potently activated by an inert ligand. Proc Natl Acad Sci U S A 104, 5163–5168 (2007). 10.1073/pnas.0700293104

31 Urban, D. J. & Roth, B. L. DREADDs (designer receptors exclusively activated by designer drugs): chemogenetic tools with therapeutic utility. Annu Rev Pharmacol Toxicol 55, 399–417 (2015). 10.1146/annurev-pharmtox-010814-124803

32 Geddes, S. D. et al. Target-specific modulation of the descending prefrontal cortex inputs to the dorsal raphe nucleus by cannabinoids. Proc Natl Acad Sci U S A 113, 5429–5434 (2016). 10.1073/pnas.1522754113

33 Holloway, B. B. et al. Monosynaptic glutamatergic activation of locus coeruleus and other lower brainstem noradrenergic neurons by the C1 cells in mice. J Neurosci 33, 18792–18805 (2013). 10.1523/JNEUROSCI.2916-13.2013

34 Kim, J. C. et al. Linking genetically defined neurons to behavior through a broadly applicable silencing allele. Neuron 63, 305–315 (2009). 10.1016/j.neuron.2009.07.010

35 Schiavo, G., Matteoli, M. & Montecucco, C. Neurotoxins affecting neuroexocytosis. Physiol Rev 80, 717–766 (2000). 10.1152/physrev.2000.80.2.717

36 Chefer, V. I., Backman, C. M., Gigante, E. D. & Shippenberg, T. S. Kappa opioid receptors on dopaminergic neurons are necessary for kappa-mediated place aversion. Neuropsychopharmacology 38, 2623–2631 (2013). 10.1038/npp.2013.171

37 Mucha, R. F. & Herz, A. Motivational properties of kappa and mu opioid receptor agonists studied with place and taste preference conditioning. Psychopharmacology (Berl) 86, 274–280 (1985). 10.1007/BF00432213

38 Ohla, K. et al. Recognizing Taste: Coding Patterns Along the Neural Axis in Mammals. Chem Senses 44, 237–247 (2019). 10.1093/chemse/bjz013

39 Oliveira-Maia, A. J., Roberts, C. D., Simon, S. A. & Nicolelis, M. A. Gustatory and reward brain circuits in the control of food intake. Adv Tech Stand Neurosurg 36, 31–59 (2011). 10.1007/978-3-7091-0179-7_3

40 Azevedo, F. A. et al. Equal numbers of neuronal and nonneuronal cells make the human brain an isometrically scaled-up primate brain. J Comp Neurol 513, 532–541 (2009). 10.1002/cne.21974

41 von Bartheld, C. S., Bahney, J. & Herculano-Houzel, S. The search for true numbers of neurons and glial cells in the human brain: A review of 150 years of cell counting. J Comp Neurol 524, 3865–3895 (2016). 10.1002/cne.24040

42 Kim, D.-I. et al. Novel genetically encoded tools for imaging or silencing neuropeptide release from presynaptic terminals *in vivo*. bioRxiv, 2023.2001.2019.524797 (2023). 10.1101/2023.01.19.524797

43 Condon, L. F. et al. Parabrachial Calca neurons drive nociplasticity. Cell Rep 43, 114057 (2024). 10.1016/j.celrep.2024.114057

44 Norgren, R. & Leonard, C. M. Taste pathways in rat brainstem. Science 173, 1136–1139 (1971). 10.1126/science.173.4002.1136

45 Krout, K. E. & Loewy, A. D. Parabrachial nucleus projections to midline and intralaminar thalamic nuclei of the rat. J Comp Neurol 428, 475–494 (2000). 10.1002/1096-9861(20001218)428:3<475::aid-cne6>3.0.co;2-9

46 Lasiter, P. S. & Kachele, D. L. Postnatal development of the parabrachial gustatory zone in rat: dendritic morphology and mitochondrial enzyme activity. Brain Res Bull 21, 79–94 (1988). 10.1016/0361-9230(88)90122-0

47 Tokita, K., Inoue, T. & Boughter, J. D., Jr. Subnuclear organization of parabrachial efferents to the thalamus, amygdala and lateral hypothalamus in C57BL/6J mice: a quantitative retrograde double labeling study. Neuroscience 171, 351–365 (2010). 10.1016/j.neuroscience.2010.08.026

48 Jarvie, B. C., Chen, J. Y., King, H. O. & Palmiter, R. D. Satb2 neurons in the parabrachial nucleus mediate taste perception. Nat Commun 12, 224 (2021). 10.1038/s41467-020-20100-8

49 Kang, S. J. et al. A central alarm system that gates multi-sensory innate threat cues to the amygdala. Cell Rep 40, 111222 (2022). 10.1016/j.celrep.2022.111222

50 Ogawa, H. & Nomura, T. Receptive field properties of thalamo-cortical taste relay neurons in the parvicellular part of the posteromedial ventral nucleus in rats. Exp Brain Res 73, 364–370 (1988). 10.1007/BF00248229

51 Nakashima, M. et al. An anterograde and retrograde tract-tracing study on the projections from the thalamic gustatory area in the rat: distribution of neurons projecting to the insular cortex and amygdaloid complex. Neurosci Res 36, 297–309 (2000). 10.1016/s0168-0102(99)00129-7

52 Yasui, Y., Itoh, K. & Mizuno, N. Projections from the parvocellular part of the posteromedial ventral nucleus of the thalamus to the lateral amygdaloid nucleus in the cat. Brain Res 292, 151–155 (1984). 10.1016/0006-8993(84)90899-0

53 Yasui, Y., Itoh, K., Sugimoto, T., Kaneko, T. & Mizuno, N. Thalamocortical and thalamo- amygdaloid projections from the parvicellular division of the posteromedial ventral nucleus in the cat. J Comp Neurol 257, 253–268 (1987). 10.1002/cne.902570210

54 Haley, M. S., Fontanini, A. & Maffei, A. Inhibitory Gating of Thalamocortical Inputs onto Rat Gustatory Insular Cortex. J Neurosci 43, 7294–7306 (2023). 10.1523/JNEUROSCI.2255-22.2023

55 Fletcher, M. L., Ogg, M. C., Lu, L., Ogg, R. J. & Boughter, J. D., Jr. Overlapping Representation of Primary Tastes in a Defined Region of the Gustatory Cortex. J Neurosci 37, 7595–7605 (2017). 10.1523/JNEUROSCI.0649-17.2017

56 Peng, Y. et al. Sweet and bitter taste in the brain of awake behaving animals. Nature 527, 512–515 (2015). 10.1038/nature15763

57 Wang, L. et al. The coding of valence and identity in the mammalian taste system. Nature 558, 127–131 (2018). 10.1038/s41586-018-0165-4

58 Blair, H. T., Schafe, G. E., Bauer, E. P., Rodrigues, S. M. & LeDoux, J. E. Synaptic plasticity in the lateral amygdala: a cellular hypothesis of fear conditioning. Learn Mem 8, 229–242 (2001). 10.1101/lm.30901

59 Maren, S. & Quirk, G. J. Neuronal signalling of fear memory. Nat Rev Neurosci 5, 844–852 (2004). 10.1038/nrn1535

60 Roman, C. W., Derkach, V. A. & Palmiter, R. D. Genetically and functionally defined NTS to PBN brain circuits mediating anorexia. Nat Commun 7, 11905 (2016). 10.1038/ncomms11905

61 Park, S., Zhu, A., Cao, F. & Palmiter, R. Parabrachial *Calca* neurons mediate second-order conditioning. bioRxiv, 2024.2003.2021.586150 (2024). 10.1101/2024.03.21.586150

62 Zhou, P. et al. Efficient and accurate extraction of in vivo calcium signals from microendoscopic video data. Elife 7 (2018). 10.7554/eLife.28728

63 Fadok, J. P., Dickerson, T. M. & Palmiter, R. D. Dopamine is necessary for cue-dependent fear conditioning. J Neurosci 29, 11089–11097 (2009). 10.1523/JNEUROSCI.1616-09.2009

